# EnsPlex: Integrative Protein Complex Prediction through Multi-source Complementary Structural Sampling and Topology-Aware Candidate Selection

**DOI:** 10.64898/2026.09.26.754739

**Authors:** Baochuan Hu, Zhenhua Lu, Khalid Zaman, Zhaoxi Sun, Xiao He

## Abstract

Protein complex structure prediction depends on both the breadth of candidate coverage and the ability to select accurate models within a limited output budget. Most existing approaches emphasize only one stage. End-to-end models and molecular docking workflows generate candidate structures, whereas quality-assessment models primarily rerank a predefined candidate pool. A complete strategy must account for complementary sampling across generators, differences among scoring scales and the allocation of final candidate quotas. Here, EnsPlex is presented as a multi-source framework that couples structural sampling with candidate selection for protein-protein interaction complex prediction. EnsPlex combines conformation-expanded docking, AlphaFold-Multimer, AlphaFold3 and Boltz-1 to expand the sampled conformational space. Conformation-expanded docking comprises monomer conformational expansion followed by flexible HADDOCK docking. FACET is trained to predict candidate quality using DockQ and its component metrics as supervision. Across 102 antigen-antibody systems, EnsPlex achieved up to a 27.8% relative improvement in target success rate over AlphaFold3 with its built-in ranking under matched output budgets. FACET also improved within-source ranking in the internal candidate pools and in homology-filtered external data. These findings support the utility of EnsPlex for structure prediction in the evaluated antigen-antibody systems.

## 1 Introduction

Protein-protein interactions (PPIs) mediate essential biological processes, including signal transduction, immune recognition, metabolic regulation and gene expression. Determining the three-dimensional structures of PPI complexes helps elucidate molecular recognition mechanisms and provides a structural basis for antibody engineering, protein design and targeted drug development. Unlike monomer structure prediction, complex structure prediction must account for component flexibility, interface location, relative orientation and overall assembly within a substantially larger conformational search space. A practically useful strategy must therefore not only generate at least one model of the desired quality but also identify it within a limited set of outputs.

Several classes of candidate-generation and docking methods have been developed. End-to-end models, including AlphaFold-Multimer, AlphaFold3 and Boltz-1, generate complex structures and associated confidence estimates directly from sequences or molecular compositions. By contrast, workflows such as HADDOCK, ClusPro and AbAdapt explore the assembly space through rigid-body searches, restrained docking and local flexible refinement^1–6^. These workflows typically start from monomer structures, interface information or homology models. However, each method generally produces its own candidate set and applies its own built-in ranking scheme, complicating comparisons across models. Moreover, different generators may fail to sample near-native regions for different targets, particularly in antigen-antibody systems characterized by weak coevolutionary signals and flexible interfaces. Consequently, candidates from a single source may not simultaneously provide broad conformational coverage and a consistently high probability of including accurate models^7, 8^.

A complementary body of work focuses on candidate-structure quality assessment. Traditional docking scoring functions rank decoys using energy terms, empirical potentials or interfacial contacts^3^. Data-driven methods, including DeepRank, DeepRank-GNN-ESM, GDockScore, ABAG-Rank and DeepRank-Ab, learn to predict candidate quality from geometric, energetic and sequence-derived features. These methods can improve model selection within predefined candidate pools^9–13^. However, they generally do not expand conformational sampling or systematically address how candidates generated by multiple sources should be integrated for a new PPI system.

Sampling and scoring should not be treated as independent components. Even when sampling reaches a near-native region, broader conformational coverage is useful only if the scoring model can reliably recognize accurate structures generated by different methods. Moreover, raw scores produced by different tools are not calibrated to a common scale and are therefore not directly comparable. An effective integrated strategy consequently requires three capabilities: obtaining complementary conformations from different sampling methods, evaluating candidates within each source using a unified quality-assessment model and constructing a final output set that preserves the complementarity of the different candidate sources.

EnsPlex (Ensemble-sourced Sampling and Selection for PPI Complex Structure Prediction) was therefore developed as a unified framework that integrates complementary structural sampling with candidate selection for PPI complex prediction. Candidates are generated using conformation-expanded docking with HADDOCK, AlphaFold-Multimer, AlphaFold3 and Boltz-1, while FACET (Fused Atomic-Contact Evaluation via Topology-informed Prediction) is employed to rerank candidates within each source based on predicted DockQ and its component metrics. Across 102 antigen-antibody targets, which represent a highly challenging category of PPIs, EnsPlex achieved up to a 27.8% relative improvement in target success rate over AlphaFold3 with its built-in ranking under matched output budgets. FACET improved within-source ranking across candidates produced by all four generators, indicating that its performance gains were not limited to a single candidate distribution. FACET also outperformed DeepRank-GNN-ESM, GDockScore and the HADDOCK score on the homology-filtered AbSet candidates and achieved higher interface recovery rates than the AlphaFold3 ranking metrics beyond the Top-3 level on the external FoldBench evaluation.

## 2 Results

### 2.1 EnsPlex Integrates Multi-Source Sampling and Candidate Selection

For a new protein complex, a single structure-prediction tool does not always generate an acceptable model. Acceptable quality was defined as DockQ ≥ 0.23, the threshold for an acceptable model in the Critical Assessment of PRediction of Interactions (CAPRI) scheme^14^. For difficult systems, combined sampling with AlphaFold-Multimer, AlphaFold3, Boltz-1 and HADDOCK can increase coverage of acceptable conformations. Generating an acceptable conformation, however, does not ensure that it will be identified. The built-in scoring functions differ among sampling methods, and their raw scores are generally not comparable. Effective selection is therefore required within each candidate source.

EnsPlex divides complex prediction into three consecutive stages: multi-source structural sampling, within-source FACET reranking and source-balanced output. Candidate structures are generated independently from the input sequences using AlphaFold-Multimer, AlphaFold3 and Boltz-1. A separate branch expands monomer conformations through short simulations and then performs HADDOCK docking. The same pretrained FACET model scores and ranks candidates within each of the four sources. A defined quota is then selected from every source to form the final output set (Figure 1).

**Figure 1.**
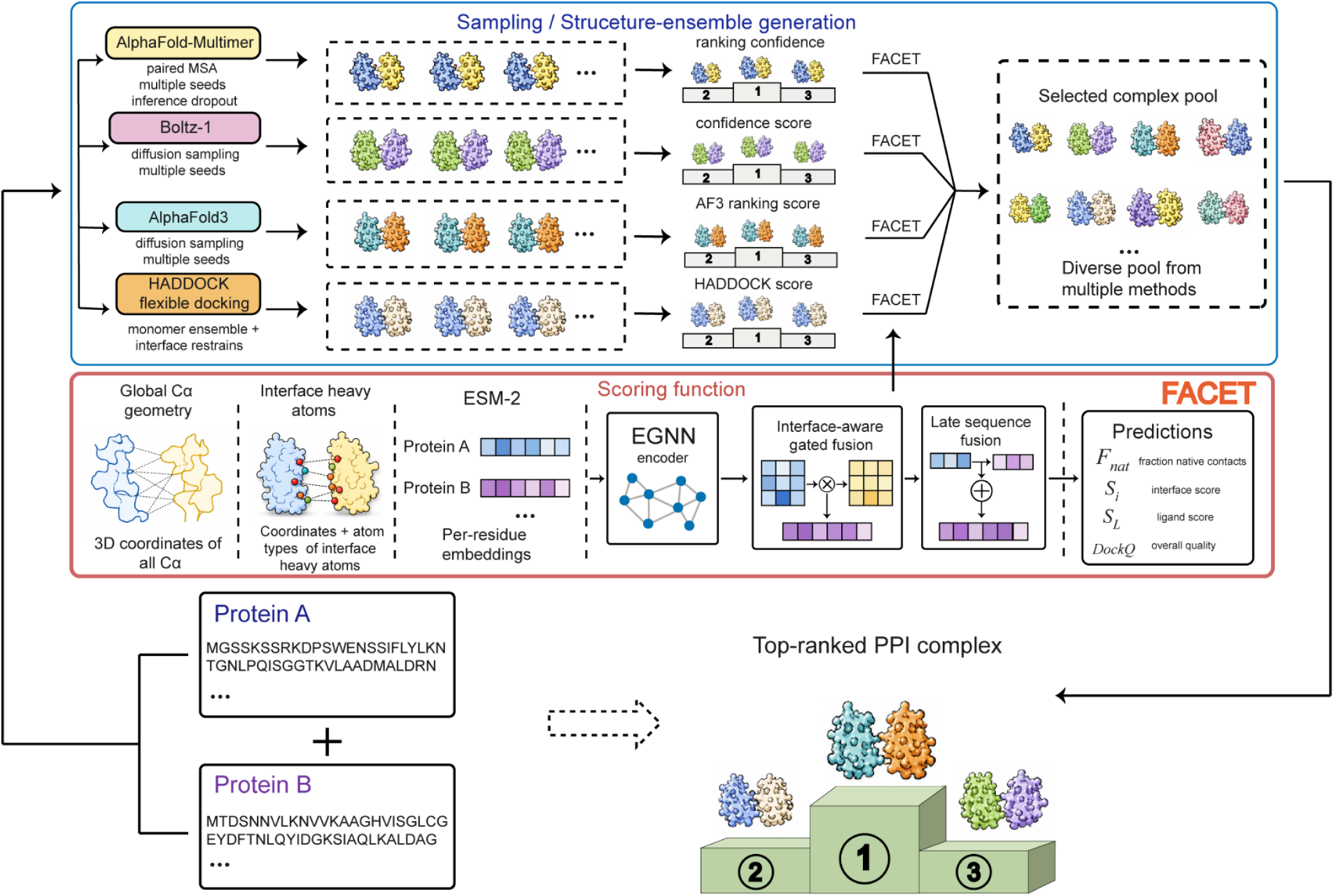
Overall framework of EnsPlex for multi-source protein complex structure prediction. Given the protein sequences of a target complex, AlphaFold-Multimer, AlphaFold3 and Boltz-1 independently generate candidate structures using multiple random seeds, while HADDOCK further expands the candidate conformational space through flexible docking with simulation-derived monomer ensembles and interface restraints. FACET evaluates candidates from all sources using a shared quality-assessment model that integrates global Cα geometry, interfacial heavy-atom features and residue-level sequence embeddings. It separately predicts Fnat, Si and SL and reconstructs DockQ from these components. FACET then reranks the candidates within each source, and the top-ranked models are retained according to a predefined source-balanced quota to form the final candidate pool and produce the prioritized complex-structure predictions.

### 2.2 Multi-Source Integration Improves Prediction Success under a Fixed Output Budget

To evaluate EnsPlex, a test set of 102 antigen–antibody targets was curated to represent the most challenging systems within PPI datasets. The complexes within the test set were first clustered and their sequence similarity was assessed with MMseqs2^15^. Complex structural similarity was also compared with Foldseek^16^. The test set was diverse and relatively balanced rather than being dominated by a few complex families (Figure S1 in the Supporting Information). Additional screening identified no obvious structurally similar antigen-antibody interfaces between the test and training sets (Figure S2 in the Supporting Information). This filtering step maintains the diversity of the test set and reduces data leakage between the training and test sets.

Together, the results in Figure 2 indicates that candidate-source complementarity and quality reranking both contributed to the prediction gains. FACET improved the ranking of acceptable models within each source, and multi-source output partly compensated for the limited conformational coverage of any single source. Figures 2A and 2B show results obtained with FACET and the original scoring functions, respectively. Figure 2C compares the complete workflow with single-source strategies using their original scores. Within a fixed output budget, EnsPlex retained the most promising structures from complementary sources.

**Figure 2.**
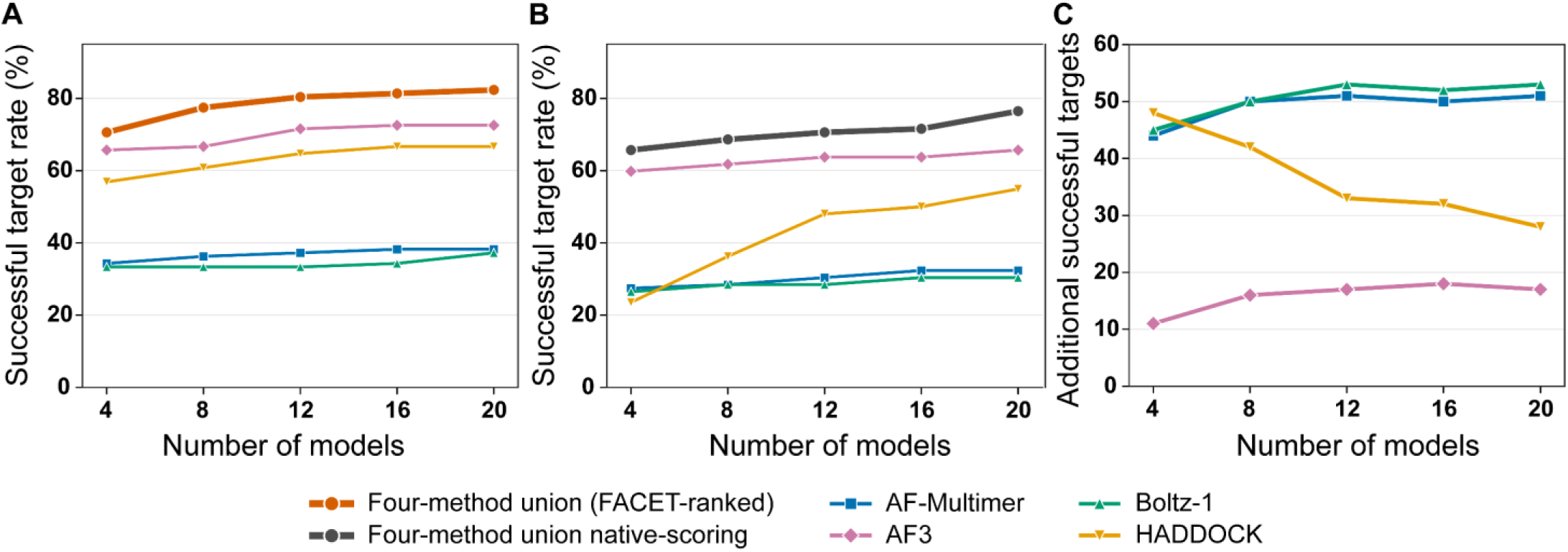
Contributions of multi-source sampling and FACET reranking under matched output budgets for 102 antigen-antibody targets. (A) Target success rates when candidates from all sources were reranked by FACET. The orange curve represents the source-balanced four-source strategy, whereas the other curves represent the corresponding single-source strategies, with candidates in each source ranked by FACET. (B) Target success rates when candidates were ranked using the original score of each generator. The gray curve represents the source-balanced four-source strategy assembled after within-source ranking with the corresponding original scores, whereas the other curves represent the original-score-ranked single-source strategies. For the four-source strategies in A and B, the total output budget was divided equally among HADDOCK, AlphaFold-Multimer, AlphaFold3 and Boltz-1. Total output budgets of 4, 8, 12, 16 and 20 correspond to retaining 1, 2, 3, 4 and 5 candidates from each source, respectively. (C) Number of additional targets successfully predicted by the FACET-ranked four-source strategy in A relative to each original-score-ranked single-source strategy in B. Each curve is calculated as the number of targets successfully predicted by the four-source FACET strategy minus that predicted by the corresponding single-source strategy indicated by the curve color. In all panels, the horizontal axis indicates the total number of structures in the final output. The vertical axis indicates the target success rate in A and B and the number of additional successful targets in C. A target was considered successfully predicted when its final output contained at least one candidate with DockQ ≥ 0.23.

### 2.3 Multi-Source Sampling Expands Candidate Coverage

To determine whether EnsPlex benefited from multi-source sampling, candidate coverage was compared across the same antigen-antibody systems. The four sources were conformation-expanded docking, AlphaFold-Multimer, AlphaFold3 and Boltz-1^1–4^. Sampling capacity was defined by the presence of a model with DockQ ≥ 0.23 in the candidate pool. For practical selection, candidates were ranked using the original score of each method. The radius of gyration (Rg) and solvent-accessible surface area (SASA) distributions of the Top-50 candidates were also compared to characterize the conformational regions explored by each generator (Figure 3).

**Figure 3.**
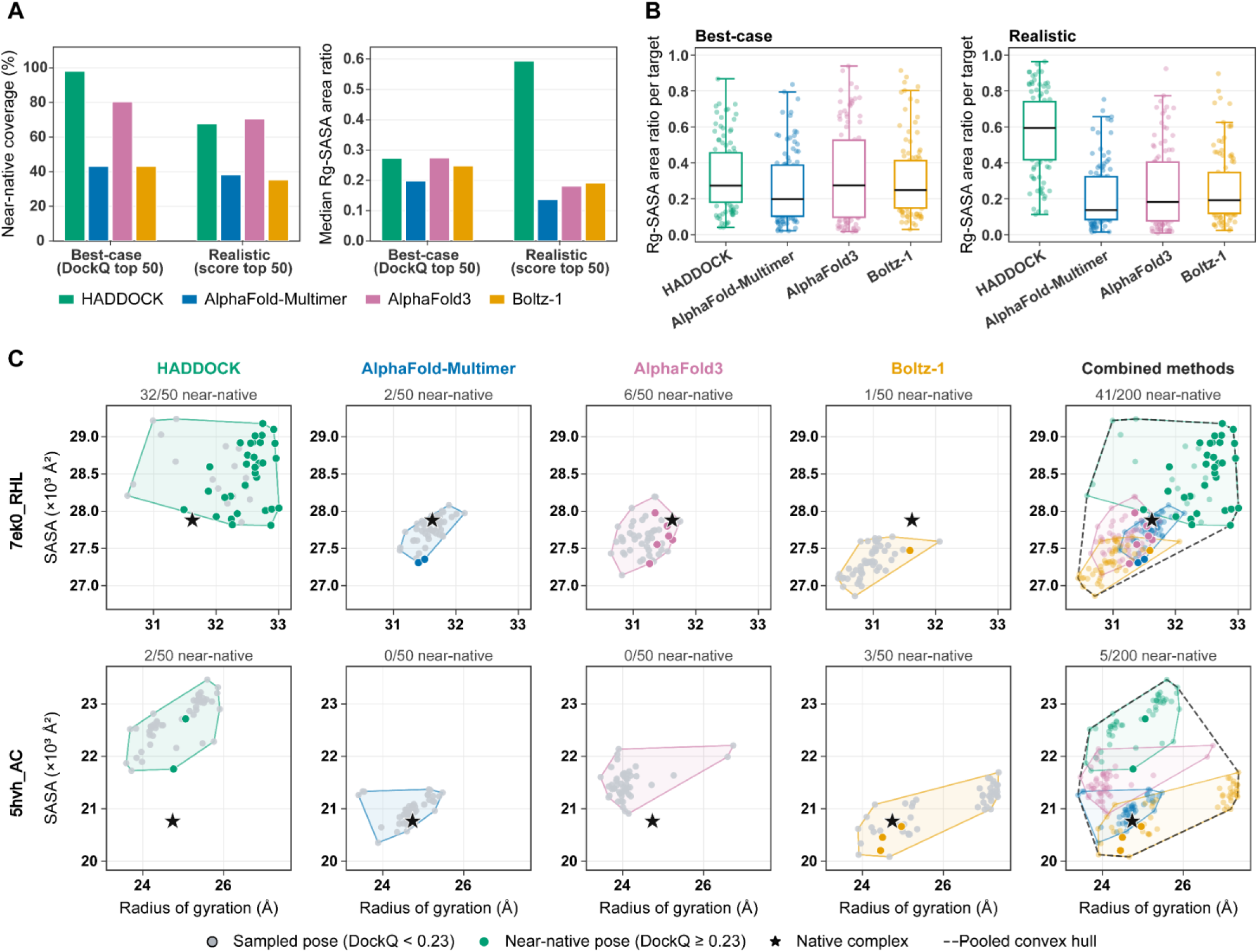
Complementary sampling coverage of the four candidate sources in *<u>Rg</u>*–SASA space. (A) Target success and *R_g_*-SASA coverage under ideal selection by DockQ and practical selection by each original score. HADDOCK, AlphaFold-Multimer, AlphaFold3 and Boltz-1 were compared on the same antigen-antibody targets using their Top-50 candidates. Left, bars show the proportion of targets with at least one selected model having DockQ ≥ 0.23. Right, the vertical axis gives the median per-target *R_g_*-SASA area ratio. A two-dimensional Gaussian kernel density estimate (KDE) was calculated from 50 models for each target and method. The area ratio equals the occupied area for one method divided by the union occupied by all four methods. A higher value indicates broader sampling. (B) Per-target *R_g_*-SASA sampling coverage for each source. The vertical axis gives the method-specific KDE area divided by the four-method union for each target. The grid and density threshold match those in panel A. Each translucent point represents one target and was calculated from 50 selected models. Boxes span the 25th to 75th percentiles, and black lines indicate medians. (C) Candidate distributions for two representative targets in *R_g_*-SASA space. For 7EK0 (top), HADDOCK sampled a broader region. For 5HVH (bottom), AlphaFold3 produced no near-native models, whereas HADDOCK and Boltz-1 did. The first four columns show HADDOCK, AlphaFold-Multimer, AlphaFold3 and Boltz-1; the fifth overlays all four methods. Each method contributed 50 models selected by its own score, yielding 200 models in each combined panel. The horizontal axis is the radius of gyration of all complex heavy atoms, *R_g_* (Å). The vertical axis is complex SASA (×10³ Å²). Panels in the same row use identical axis ranges. Gray points denote models with DockQ < 0.23, and filled colored points with white outlines denote acceptable models. Black stars mark the experimental native complexes. Numbers above each panel give near-native models divided by selected models. Translucent colored polygons are convex hulls for each method’s 50 models. The black dashed polygon is the joint convex hull of all 200 models and indicates the sampled range.

When candidates were selected by ground-truth DockQ, conformation-expanded docking achieved a 22.0% relative increase in the Top-50 target success rate over AlphaFold3. Its candidates covered a broader region of *Rg*-SASA space. However, ranking by the HADDOCK score markedly reduced the practical detection rate of acceptable models (Figures 3A and 3B). Thus, the original score did not fully identify the acceptable models already present in the candidate pool. Candidate landscapes for two representative targets showed distinct distributions across the four sources (Figure 3C). For some systems, one source reached regions that the others did not sample. These results support retaining candidates from all four sources in EnsPlex, as the different generators sampled acceptable models for complementary subsets of targets.

### 2.4 FACET Improves Within-Source Ranking and Retains Performance on External Test Sets

Beyond expanding conformational coverage, the performance of EnsPlex depends on FACET’s ability to identify candidates with the correct interface location, relative orientation and overall assembly. To isolate the effect of reranking, FACET was compared with the source-specific original scoring functions while holding each candidate pool fixed. Within the conformation-expanded docking pool, FACET achieved higher Top-k target success rates than the HADDOCK score, the HADDOCK van der Waals energy term and PRODIGY ΔG for every k from 1 to 9 (Figure 4A)^17^. FACET likewise yielded higher Top-k target success rates than the corresponding built-in ranking or confidence metric in the AlphaFold-Multimer, AlphaFold3 and Boltz-1 pools (Figures 4B–4D). Because the candidate pools were unchanged, these comparisons attribute the improvement to reranking rather than to differences in candidate generation.

**Figure 4.**
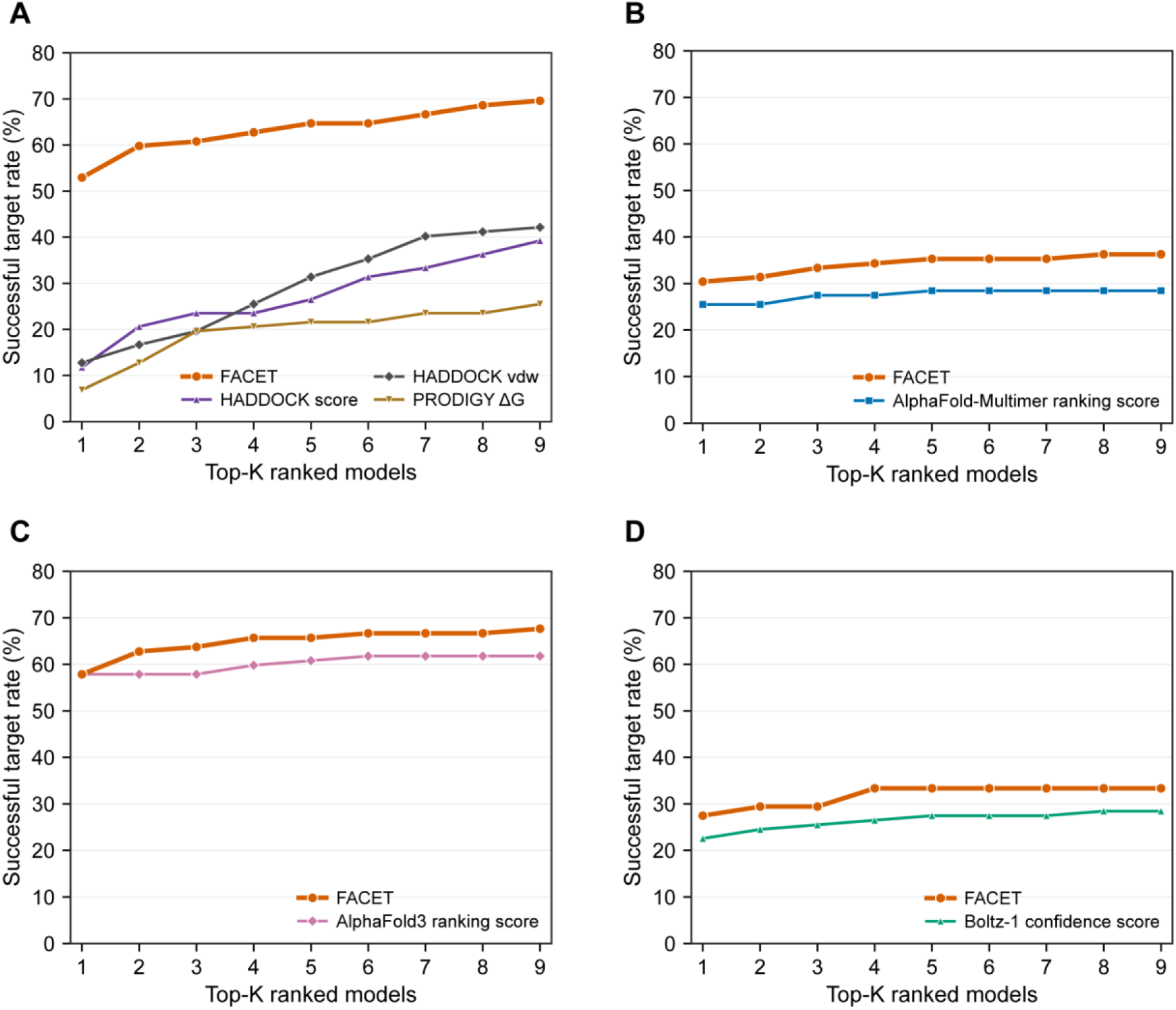
Within-source candidate-ranking performance of FACET on the internal antigen-antibody test set comprising 102 targets. (A) Top-k target success rates, for every k from 1 to 9, obtained using FACET, the HADDOCK score, the HADDOCK van der Waals energy term and PRODIGY ΔG to rank candidates from conformation-expanded docking. (B-D) Top-k target success rates obtained using FACET and the corresponding built-in ranking or confidence metric in the AlphaFold-Multimer (B), AlphaFold3 (C) and Boltz-1 (D) candidate pools. A target was considered successfully predicted at Top-k when the corresponding set contained at least one candidate with DockQ ≥ 0.23.

FACET also retained its ranking performance on external candidate pools. AbSet comprises an Ab-blind subset generated without interface restraints and an Ab-site subset generated by site-directed docking^18^. The Ab-site subset uses antibody structures predicted by ABodyBuilder2 and antigen epitope restraints derived from changes in antigen solvent-accessible surface area between the bound and unbound structures^18, 19^. To reduce overlap with the FACET training data, systems sharing more than 20% antigen sequence identity or more than 95% antibody sequence identity with any training system were removed. After filtering, the Ab-blind subset contained 201 systems and 86,500 conformations, whereas the Ab-site subset contained 127 systems and 39,250 conformations. Across the evaluated k values, FACET achieved higher Top-k target success rates than DeepRank-GNN-ESM, GDockScore and the HADDOCK score in both candidate distributions (Figures 5A and 5B)^3, 10, 11^. FACET was further evaluated on the antigen-antibody subset of FoldBench, which contains more recently released and challenging systems. After standardizing reference-chain assignments and generating candidates with AlphaFold3, 165 evaluable interfaces were identified according to the FoldBench interface definition. FACET achieved higher interface success rates than both the AlphaFold3 ranking score and chain-pair ipTM at k=4 to k=9 (Figures 5C and 5D)^20^.

**Figure 5.**
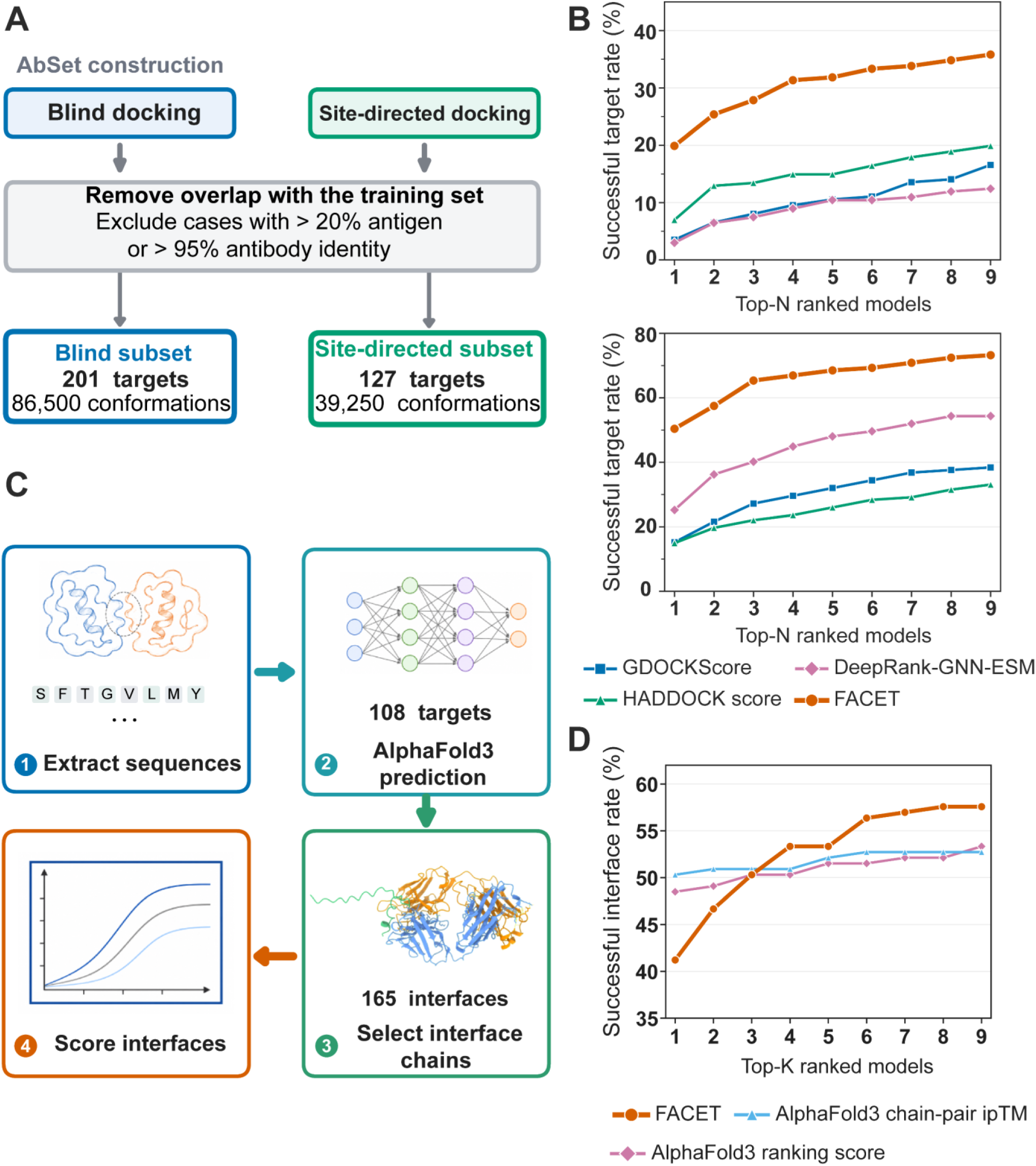
External evaluation of FACET on antigen-antibody data sets. (A) Construction of the AbSet blind-docking and site-directed subsets and their homology filtering against the FACET training set. (B) Top-k target success rates for FACET, GDockScore, DeepRank-GNN-ESM and the HADDOCK score on the filtered blind-docking subset (left) and site-directed subset (right). (C) FoldBench evaluation workflow, including sequence extraction, AlphaFold3 candidate generation, selection of reference interfaces and candidate scoring. (D) Top-k interface success rates of FACET, the AlphaFold3 ranking score and chain-pair ipTM across 165 FoldBench interfaces.

The representative cases illustrate the structural differences between candidates prioritized by FACET and those prioritized by the original scores. For 7TXT, AlphaFold3 generated an acceptable candidate, but its built-in ranking score instead prioritized a model with locally plausible interfacial contacts at the wrong binding surface. FACET assigned scores of 0.41 and 0.06 to the correctly and incorrectly bound candidates, respectively. For 8WTD, the HADDOCK score prioritized a candidate with a locally favorable interface but an incorrect overall orientation. FACET assigned scores of 0.57 and 0.04 to the correctly and incorrectly oriented candidates, respectively (Figure 6). These examples show that locally plausible interfaces can coexist with incorrect global assemblies and that FACET was able to distinguish between them. Together, the internal and external evaluations support the use of FACET as a common quality-assessment model for candidates generated by different sources.

**Figure 6.**
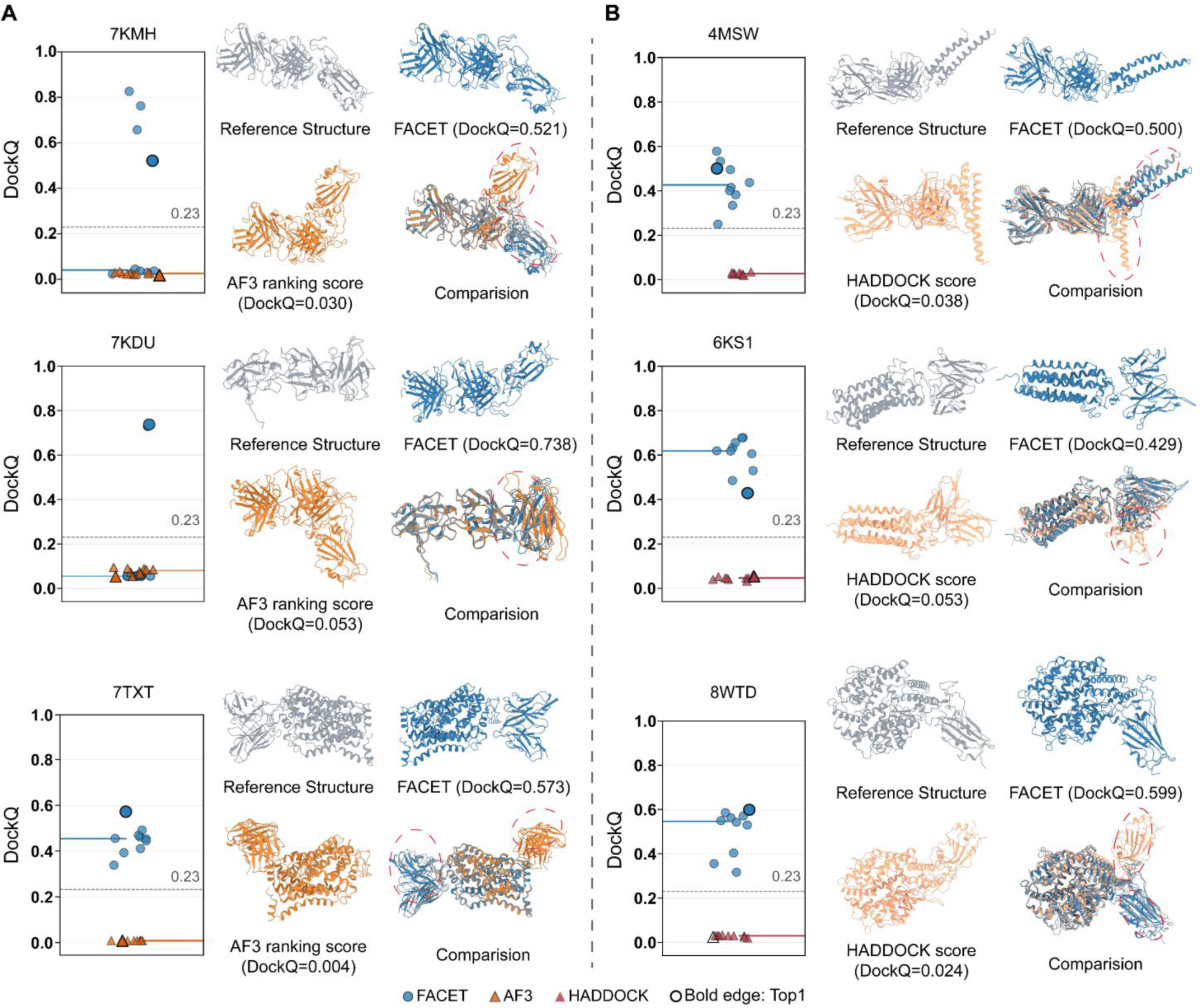
Representative structural comparisons between candidates prioritized by FACET and by the source-specific original scores. (A) AlphaFold3 candidates for 7KMH, 7KDU and 7TXT. (B) HADDOCK candidates for 4MSW, 6KS1 and 8WTD. Each panel shows the DockQ distribution, reference structure, FACET-selected model, original-score-selected model and structural superposition. The dashed line marks DockQ = 0.23, the CAPRI threshold for acceptable quality. Distribution plots show DockQ values for the Top-10 models selected by the two scoring functions. Blue circles denote FACET selections, and red triangles denote selections by the built-in score. The Top-1 selection has a thicker outline. Blue and red lines indicate the median DockQ of the respective selections. Red circles in the structural comparisons highlight major differences between the models selected by FACET and by the built-in score.

### 2.5 Scale and Diversity of the FACET Training Data

FACET was trained on a large docking dataset stratified by candidate quality and was subsequently evaluated on candidates generated by multiple methods. Starting from more than 6,000 PPI systems, over 10 million candidate structures were generated through monomer conformational expansion and flexible docking. Each candidate was annotated with DockQ and its component metrics, and these annotations were used to stratify the dataset across quality levels ranging from incorrect to high-quality models. To characterize sequence diversity and redundancy, the numbers of MMseqs2 clusters were compared at sequence identity thresholds of 50%, 70%, 90% and 100% across the FACET training set and five reference datasets: AbSet, PPI4DOCK, BM5, ABAG and the Antibody Benchmark (Figure 7A)^15, 18, 21–24^. In a t-distributed stochastic neighbor embedding (t-SNE) visualization of sequence embeddings from the ESM-2 650M model, the FACET training data were broadly distributed and extended into regions sparsely represented by several reference datasets (Figure 7B)^25^. Together, these analyses characterize the scale and sequence diversity of the training data, while the quality annotations provide supervision across a broad range of candidate accuracies. Separately, the source-species composition of the 102 antigen-antibody test targets was summarized. The test set included targets from multiple species and taxonomic groups, with their proportions and target counts shown in Figure 7C.

**Figure 7.**
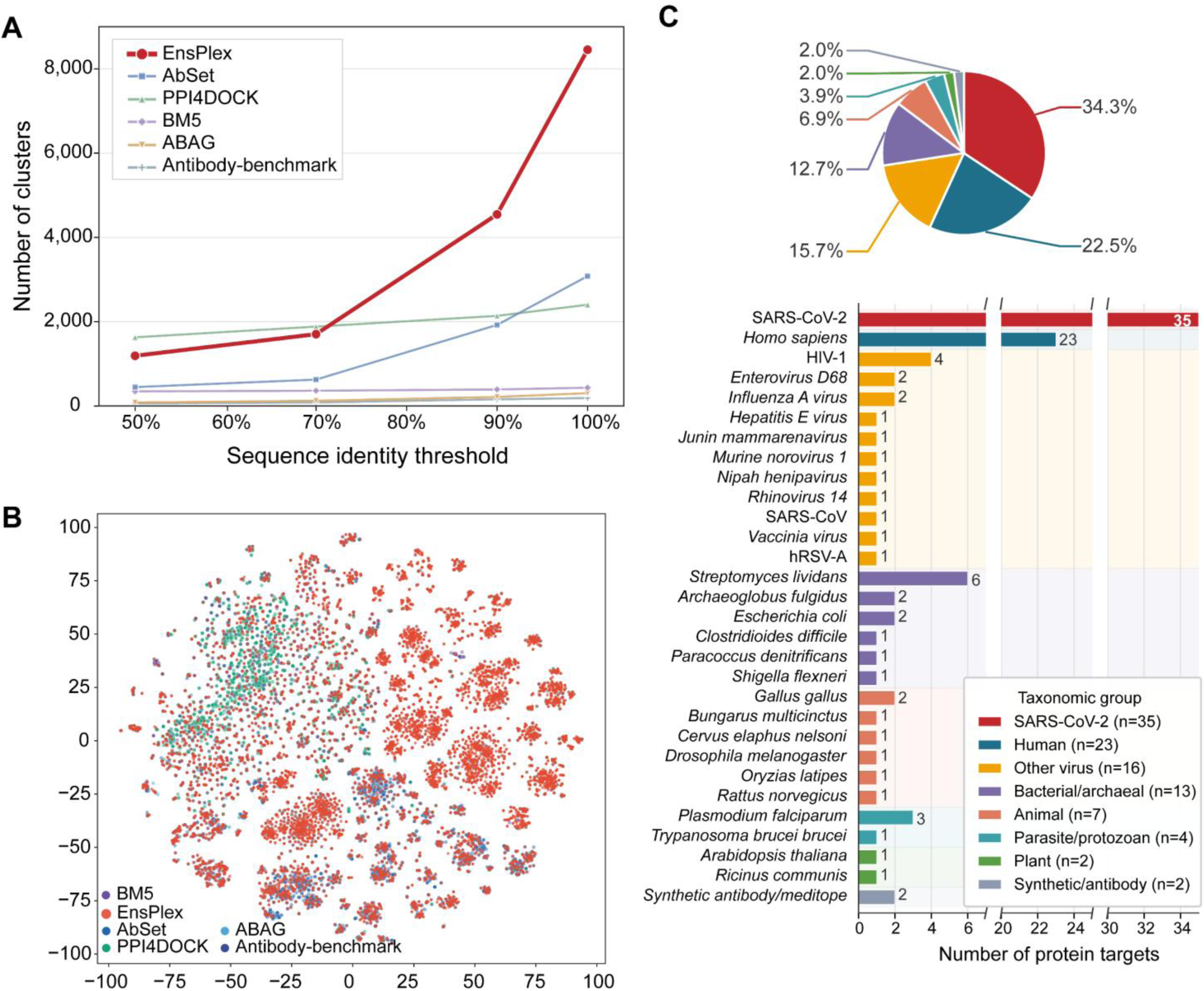
Scale and diversity of the FACET training data. (A) Numbers of sequence clusters obtained using MMseqs2 at sequence identity thresholds of 50%, 70%, 90% and 100% for the FACET training set and five reference datasets: AbSet, PPI4DOCK, BM5, ABAG and the Antibody Benchmark. The horizontal axis indicates the sequence identity threshold, and the vertical axis indicates the number of clusters. (B) t-SNE visualization of sequence embeddings generated by the ESM-2 650M model. Red points represent the FACET training data, and the other colors represent the reference datasets, as indicated in the legend. (C) Source-species composition of the 102 antigen-antibody test targets. The upper stacked bar shows the proportions of the source categories indicated in the legend, and the lower horizontal bars show the number of targets assigned to each species or category.

## 3 Discussion

EnsPlex improves antigen-antibody complex prediction by combining complementary candidate generation with effective selection under a limited output budget. Across 102 targets, the integrated workflow achieved up to a 27.8% relative improvement in target success rate over AlphaFold3 with its built-in ranking under matched output budgets. The sampling analyses and within-source comparisons suggest that this gain reflects both the recovery of acceptable structures from complementary sources and improved retention of these structures through FACET reranking. In the test set, broader sampling was useful only when the additional acceptable models were retained after reranking. Therefore, sampling coverage alone did not predict the quality of the final output.

The distinction between generating acceptable structures and identifying them has long been recognized in protein-protein docking. Systematic evaluations have shown that scoring performance depends on docking difficulty and candidate quality, while CAPRI scoring benchmarks assess selection independently of candidate generation using models from multiple prediction methods^26, 27^. Antigen-antibody prediction presents additional challenges. Antibody sequence diversity arises through V(D)J recombination and somatic hypermutation,^28^ while weak interchain coevolutionary signals and uncertainty in complementarity-determining-region conformations complicate the prediction of binding sites and relative orientations^8, 20^. Consequently, plausible local contacts or high internal confidence do not necessarily indicate a correct assembly. This motivates evaluating candidate quality using both interfacial information and the broader structural context.

Previous studies have addressed these limitations through expanded sampling and improved scoring. AFsample demonstrated the benefits of extensive sampling with varied inference settings, while antigen-antibody benchmarks revealed a persistent gap between the best generated structure and the model selected by internal confidence scores^8, 29^. AlphaFold2-based refinement and rescoring also improved the selection of docking-generated models, although combining multiple docking methods yielded only modest additional improvements at the top of the ranked list^30^. In these evaluations, individual candidate sources recovered acceptable structures for targets missed by others, supporting their complementary use. The gap between DockQ-based selection and HADDOCK-score-based selection further showed that useful conformations could remain overlooked within the docking pool. EnsPlex addresses both issues by combining heterogeneous generators with FACET-based selection within each source.

Learning-based methods such as DeepRank-GNN-ESM and GDockScore have demonstrated the value of graph-based representations for candidate-quality assessment^10, 11^. FACET combines global complex geometry, interfacial atomic contacts and sequence information to assess local interfaces in the context of the overall assembly. Its representative selections illustrate why this distinction matters: locally plausible interfaces can occur at the wrong binding surface or in an incorrect orientation. DeepRank-Ab and ABAG-Rank have also investigated transfer across candidate-generation methods, highlighting the importance of evaluating performance under changes in candidate distribution^12, 13^. Emphasis is placed on within-target selection performance: FACET improved ranking in fixed internal candidate pools and retained ranking gains on the filtered AbSet pools and AlphaFold3-generated FoldBench candidates. FACET therefore transferred to the four candidate sources tested here, as well as to the filtered AbSet and AlphaFold3-generated FoldBench candidates.

The present evaluation leaves several practical questions open. The number of final predictions, rather than total computational cost, was matched, so the benefit of EnsPlex under a fixed compute budget remains to be tested. The HADDOCK branch also used interface restraints, meaning that its contribution partly depends on the availability and quality of prior interface information. In addition, the final pool was assembled using equal quotas from the four sources; whether target-specific allocation or cross-source score calibration could achieve similar or better coverage with fewer structures remains unclear. Testing additional antibody formats, antigen classes and other PPI systems is needed to determine how broadly the observed benefit extends beyond the present benchmark.

## 4 Conclusion

Protein complex structure prediction depends on both generating acceptable conformations and identifying them among a limited number of reported models. EnsPlex addresses these coupled challenges by combining conformation-expanded docking, AlphaFold-Multimer, AlphaFold3 and Boltz-1 with FACET-based within-source ranking and source-balanced selection. Across 102 antigen–antibody targets, EnsPlex improved target success by up to 27.8% relative to AlphaFold3 with its native ranking under matched output budgets. Comparisons within fixed candidate pools showed that FACET improved the selection of acceptable models, while the sampling analyses indicated that the four generators provided partly complementary target coverage. FACET also retained ranking gains on the homology-filtered AbSet candidates and in the external FoldBench evaluation. In conclusion, these findings support integrating complementary sampling with effective candidate selection, rather than improving either step in isolation.

The present evidence is limited to the evaluated antigen–antibody systems and matched output counts; the benefit under matched computational costs remains unknown, and the docking branch depends on available interface information. Future work should examine compute-matched performance, alternative source-allocation strategies and applicability to other PPI classes.

## 5 Methods and Data Preparation

### 5.1 Training Candidates and Conformation-Expanded Docking

FACET training candidates were generated by monomer conformational expansion followed by flexible HADDOCK docking. The same workflow provides a complementary sampling branch when EnsPlex predicts a new system. For more than 6,000 PPI systems, AlphaFold 2.3.1 first predicted the structure of each monomeric component. The model with the highest confidence was selected as the starting conformation, with antibody heavy and light chains treated as one component^31^. Short all-atom molecular dynamics simulations were then performed with the AMBER ff14SB force field and explicit solvent to expand each monomer ensemble^32^.

Complexes were assembled by guided flexible docking with HADDOCK. Twelve conformational templates were retained for both antibody and antigen, producing 12 × 12 template combinations. Ambiguous interaction restraints were applied at the interface to represent epitope information^3^. All candidates were labeled with DockQ and its Fnat, Si and SL components. Quality stratification and distribution control were then used to construct the training set. The data-generation workflow is shown in Figure S3 in the Supporting Information.

### 5.2 FACET Candidate-Quality Model and Training

FACET treats docking-quality assessment as supervised regression. For each two-chain candidate, DockQ is defined as the mean of three components^14^.

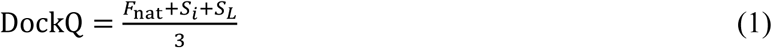

Here, *F_nat_* measures recovered native residue contacts, whereas Si and SL map the iRMSD and LRMSD (Å) to [0, 1]:

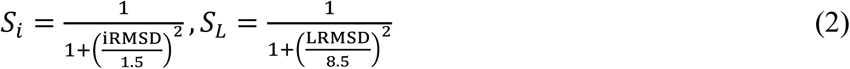

FACET predicts Fnat, Si and SL separately and reconstructs DockQ, providing complementary supervision for native contacts and overall conformation. Its structure-first architecture combines global Cα and interfacial heavy-atom encoders, gated structural fusion, late ESM-2 sequence fusion and adaptive multi-branch readout (Figure 8).

**Figure 8.**
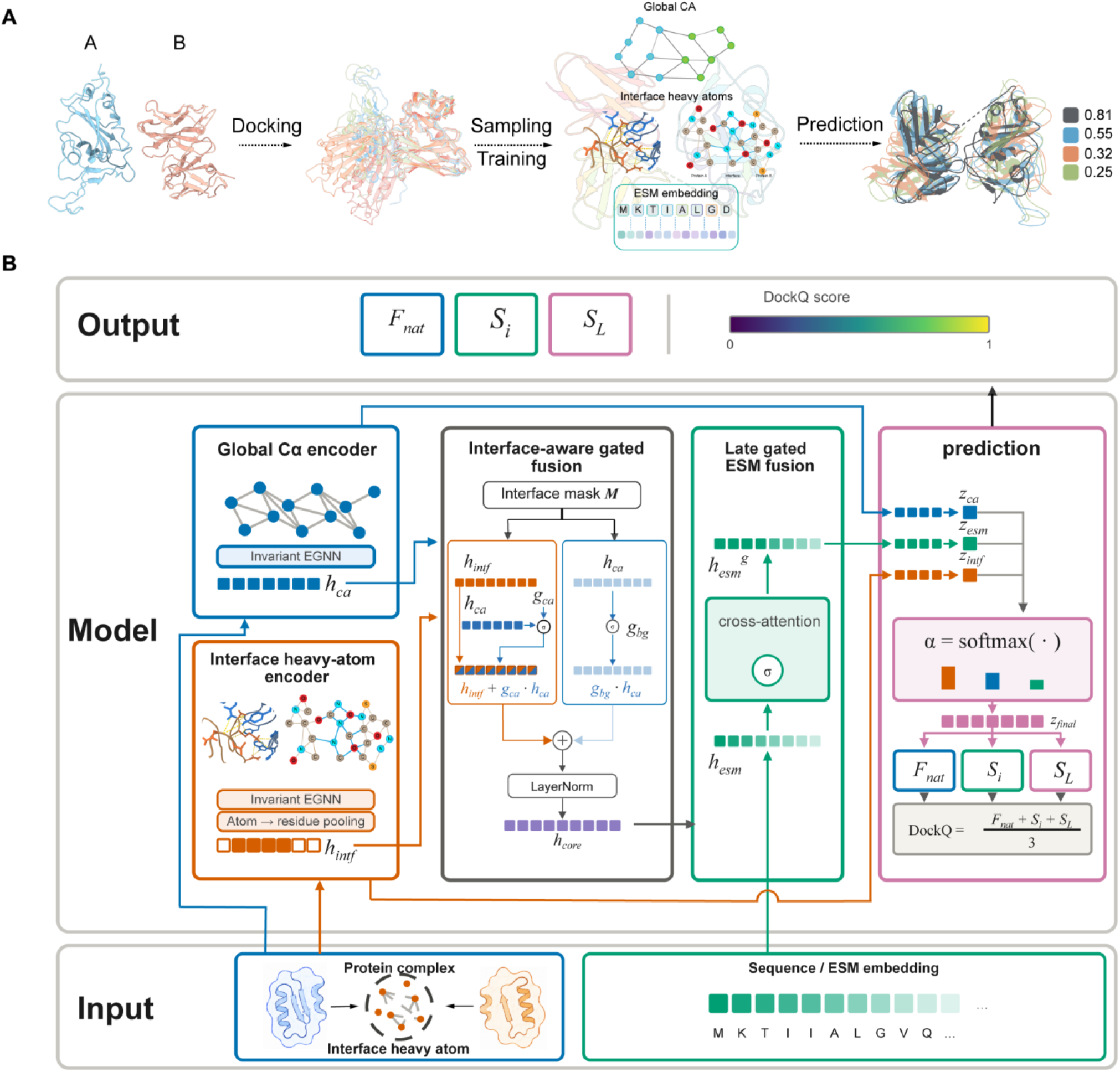
FACET candidate-quality model and training objectives. (A) Monomer conformational expansion and flexible HADDOCK docking generate quality-stratified candidates supervised by *Fnat, Si, SL and DockQ. (B) The model combines* global Cα encoding, interfacial heavy-atom encoding, interface-aware gating and late sequence fusion. It predicts Fnat, Si and SL and reconstructs DockQ.

### 5.3 Input Representations and Model Architecture

#### Input representations

Each standardized complex is represented at residue and interface-atom resolutions. Residue features comprise amino acid type, chain and intrachain indices, Cα coordinates, a local backbone frame, the Cα-to-Cβ vector and optional ESM-2 embeddings. Interface-atom features include coordinates, atom type, residue and chain membership, backbone or side-chain identity and validity masks. Edge features use invariant distance expansions, local-frame displacements and rotations, as well as interchain and sequence indicators (Sections S4.2-S4.5 in the Supporting Information).

#### Invariant message-passing layer

Both encoders use the same invariant kernel, which updates node features without changing coordinates. For node *i* with neighborhood *N*(*i*), one layer applies:

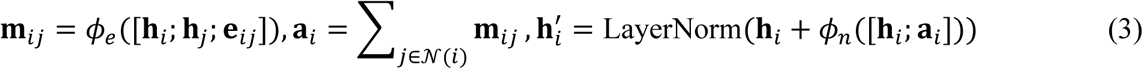

Here, *e_ij_* denotes invariant edge features, and *φ_e_* and *φ_n_* are multilayer perceptrons.

#### Global Cα residue-graph encoder

A k-nearest-neighbor graph is built from all Cα atoms within each complex. Node and edge features encode residue identity, local backbone geometry, Cα distances, sequence separation and chain relationships. Linear projection followed by message passing produces the global-context embedding (Section S4.3 in the Supporting Information).

#### Interfacial heavy-atom graph encoder

Interface residues are defined by a 6 Å interchain heavy-atom cutoff, and a distance-threshold graph is built over their heavy atoms. Features encode atom and residue identity, interatomic distances and same-residue or interchain relationships. Attention then pools atom-level features to residue representations, with noninterface positions set to zero (Sections S4.4-S4.5 in the Supporting Information).

#### Interface-dominant structural fusion

After independent linear projections align the two branches, position-specific gates fuse interface and global representations. Interface and noninterface residues are processed separately. For interface residues, the atomic representation remains dominant while a gate adds global context:

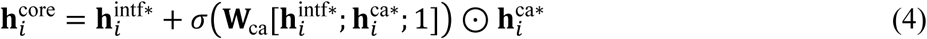

For noninterface residues, only a gated global background is retained:

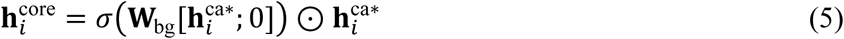

A negative gate bias keeps interface features dominant early in training. The layer-normalized fused representation *h^core^* then forms the query for sequence fusion.

#### Late gated sequence fusion

Protein-language-model features can support PPI modeling. FACET projects ESM-2 embeddings to 128 dimensions^25^ and introduces them after structural fusion through two gated cross-attention blocks. Interface-residue representations act as queries, while sequence embeddings from all residues provide keys and values:

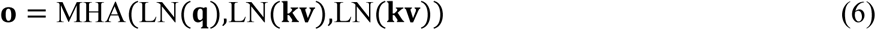

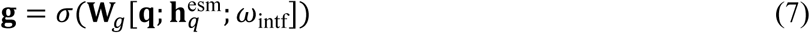

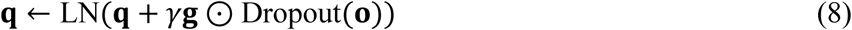

Here, *ω_intf_* is the soft interface weight and *γ* the channel-wise LayerScale factor. Gate-bias and LayerScale initialization restrict early sequence updates, preserving geometry as the primary signal.

#### Dynamic multi-branch readout and component-level prediction

The interface, global and sequence-fused branches are pooled separately over interface residues by attention:

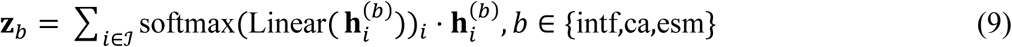

A lightweight network assigns sample-dependent fusion weights:

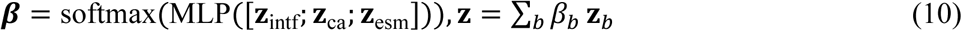

The fused vector *z* enters a shared projection and three prediction heads:

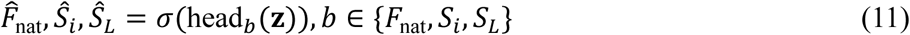

Sigmoid activations constrain all outputs to [0, 1], and DockQ is reconstructed as:

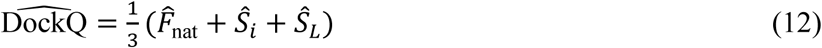

## Supporting information

support information

## Acknowledgments

This work was supported by the National Natural Science Foundation of China (Grant Nos. 62305285, 22673072, 22633005, 92477103 and 22273023), Shanghai Municipal Science and Technology Commission (Grant Nos. 25511102400 and 25DX2800500), Shanghai Municipal Commission of Economy and Informatization with Grant No. 2026-GZL-RGZN-01015, Advanced Materials-National Science and Technology Major Project (Grant No. 2026ZD0623702), the Shanghai Frontiers Science Center of Molecule Intelligent Syntheses and the Fundamental Research Funds for the Central Universities. The authors also acknowledge the Supercomputer Center of East China Normal University (ECNU Multifunctional Platform for Innovation 001) for providing computer resources.

## Author Contributions

Baochuan Hu: Data curation, Formal analysis, Investigation, Methodology, Software, Validation, Visualization, Writing - original draft, Writing - review & editing.

Zhenhua Lu: Data curation, Formal analysis, Investigation, Methodology, Software, Validation, Visualization, Writing - original draft, Writing - review & editing.

Khalid Zaman: Formal analysis, Investigation, Validation, Writing - original draft, Writing - review & editing.

Zhaoxi Sun: Conceptualization, Data curation, Formal analysis, Funding acquisition, Investigation, Methodology, Project administration, Resources, Software, Supervision, Validation, Visualization, Writing - original draft, Writing - review & editing.

Xiao He: Conceptualization, Funding acquisition, Investigation, Methodology, Project administration, Resources, Supervision, Validation, Visualization, Writing - original draft, Writing - review & editing.

## Competing interests

The authors declare no conflict of interest.

## Data Availability

The EnsPlex source code is available at https://github.com/H18020306/EnsPlex, and the trained model weights for EnsPlex are available at https://doi.org/10.5281/zenodo.22858234. The reference PDB files used for training and evaluation are available at https://doi.org/10.5281/zenodo.22857353.

## Supporting Materials

Additional analyses of complex-level homology within the 102-target antigen–antibody test set and interface similarity between the test and training sets; details of conformation-expanded docking, dataset composition and training-candidate quality coverage; FACET input representations, architecture and training objectives; ablation and graph-neural-network baseline evaluations; and representative candidate-reranking cases (PDF).

## Notes

https://doi.org/10.5281/zenodo.22857353

https://doi.org/10.5281/zenodo.22858234

https://github.com/H18020306/EnsPlex

### Competing Interest Statement

The authors have declared no competing interest.

