## Supplementary material for "EnsPlex: Integrative Protein Complex Prediction through Multi-source Complementary Structural Sampling and Topology-Aware Candidate Selection": support information

### S1. Complex clustering of the internal EnsPlex antigen–antibody test set

We performed a complex-level analysis of the antigen-antibody systems in the test set. The 102 targets showed substantial diversity at the complex level, and the scoring function retains its advantage after retaining one representative per homology component (Supplementary Fig. S1).

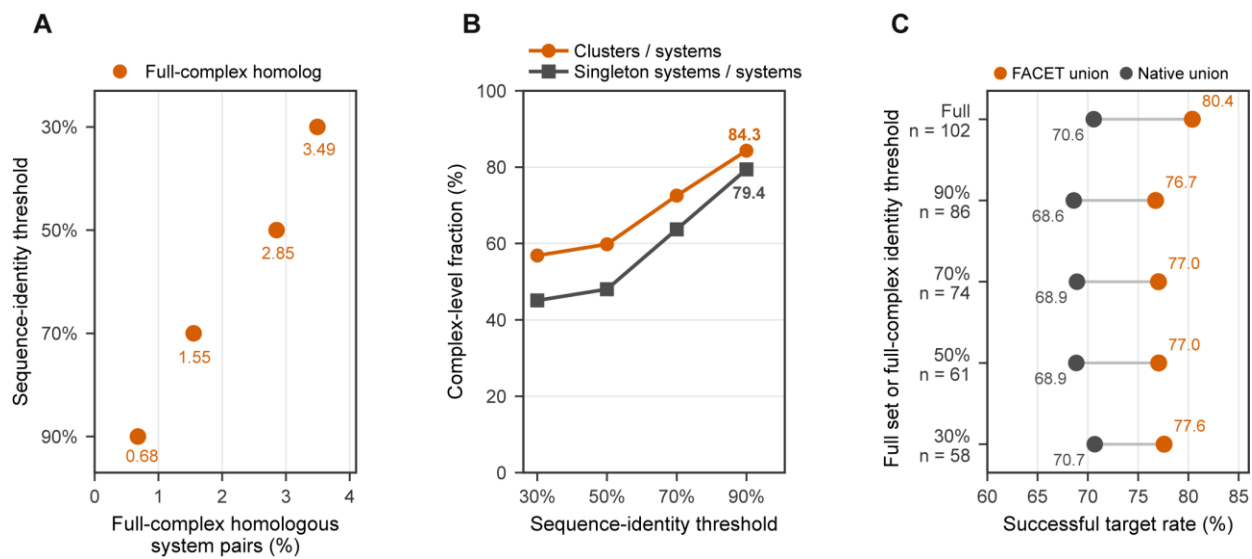

**Figure S1. Test-set homology, complex-level sequence diversity, and sampling performance after homology deduplication.** (A) Percentage of 5,151 unordered test-system pairs classified as full-complex homologs at MMseqs2 sequence-identity thresholds of 30%, 50%, 70%, and 90%; the corresponding values are 3.49%, 2.85%, 1.55%, and 0.68%. A full-complex homolog requires one-to-one matches for all chains with at least 80% query and target coverage. (B) Cluster richness (orange) and singleton-system fraction (gray) among 102 complexes at the same thresholds; clusters are connected components of the full-complex homology graph. (C) Successful-target rates at a fixed output budget of 12 after retaining one representative per homology component. The full set contains 102 targets; the 90%, 70%, 50%, and 30% subsets contain 86, 74, 61, and 58 targets. Orange and gray indicate the FACET-ranked and native-score four-source unions, respectively. Success required DockQ $\geq 0.23$.

### S2. Interface similarity between the internal EnsPlex antigen-antibody test set and the training set

For test-set systems, we used Foldseek to assess antigen-antibody interface structural similarity between the test and training sets. Relevant interfaces were identified according to the antigen–antibody chain composition of each system and compared with a common interface representation using Foldseek, where the sensitivity was set to 9.5 and the E-value cutoff was $1e-3$. No interface hit records were identified for the 102 test targets in the training-set search results (Supplementary Fig. S2).

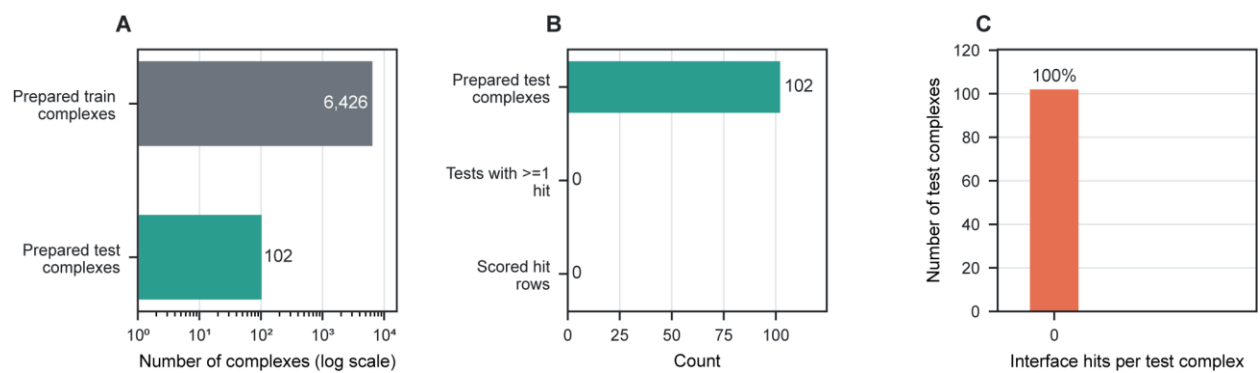

**Figure S2. Foldseek analysis of antigen-antibody chain-pair interfaces between the internal test and training sets.** (A) Input sizes for the 102 test complexes and 6,426 training complexes; the x axis is logarithmic. (B) Search funnel from prepared test complexes to interfaces with at least one hit and scored interface rows. No test complex produced a hit or score record. (C) Interface-hit counts per test complex; all 102 targets had zero hits. Zeros indicate absent search output rather than an interface TM-score of zero, and no interface-difference value was assigned.

#### S3. Conformation-Augmented Docking: Data Generation and Quality Coverage

For each protein monomer, we selected the highest-confidence AlphaFold model as the initial structure and generated a monomer conformational ensemble by short all-atom molecular-dynamics (MD) simulations in explicit solvent.

##### S3.1 Constraint-guided HADDOCK docking

For each protein complex, we constructed residue-contact restraints to simulate epitope guidance. These restraints were defined as simulated interface priors to broaden conformational sampling and do not represent experimentally measured epitopes or the native interface. Candidate interface residues on the antibody were kept fixed, whereas active residues on the antigen or receptor-binding domain (RBD) were varied between docking runs. Different restraint templates broadened the sampled conformations and increased the chance of obtaining structures close to the experimental complex. Docking was performed with HADDOCK 2.4 (2022-01 release). Ambiguous interaction restraints (AIRs) introduced this simulated antigen/RBD guidance during conformational sampling; only the selected antigen/RBD residues differed between runs.

##### S3.2 HADDOCK scoring functions

The structures were ranked with the following weighted energy functions in the three docking stages:

$$\text{Score}_{\text{it0}} = 0.01E_{\text{vdw}} + 1.0E_{\text{elec}} + 0.01E_{\text{dist}} + 1.0E_{\text{desolv}} - 0.01\text{BSA} \quad (\text{S1})$$

$$\text{Score}_{\text{it1}} = 1.0E_{\text{vdw}} + 1.0E_{\text{elec}} + 0.1E_{\text{dist}} + 1.0E_{\text{desolv}} - 0.01\text{BSA} \quad (\text{S2})$$

$$\text{Score}_{\text{refinement}} = 1.0E_{\text{vdw}} + 0.2E_{\text{elec}} + 0.1E_{\text{dist}} + 1.0E_{\text{desolv}} \quad (\text{S3})$$

#### S3.3 Conformation-augmented docking workflow and representative analysis

For each component, the highest-confidence AlphaFold monomer model was selected as the initial conformation, and short all-atom molecular-dynamics simulations in explicit solvent were used to expand conformational diversity. Complex assembly used a $12 \times 12$ flexible-docking design, combining 12 antibody templates with 12 antigen templates to produce 144 template pairs. Flexible docking was performed with the MD-expanded monomer ensembles.

AIRs were applied to the protein-protein interface in HADDOCK. Specifically, distance restraint potentials were introduced between Cα atoms of the two components. Supplementary Fig. S3A outlines the conformation-generation workflow, and Supplementary Fig. S3B shows the 7BEM antibody-antigen interface and anchor residues. Green residues denote the antigen surface, red residues denote candidate antibody surface regions and blue residues denote antigen anchor residues specified from simulated epitope information; they do not represent experimentally measured epitopes. The upper structure contains two antigen anchor residues, and the lower structure adds two further anchor points.

We used the complex formed by the SARS-CoV-2 spike receptor-binding domain and an antibody (PDB 7BEM) as a representative system to evaluate sampling by the conformation-augmented docking workflow. AlphaFold-predicted monomers were sampled with explicit-solvent molecular dynamics under the AMBER ff14SB force field to generate low-energy conformational ensembles. The radius of gyration (Rg) and solvent-accessible surface area (SASA) were used to quantify conformational differences. Supplementary Fig. S3C-F shows the distributions of the expanded antigen, antibody and assembled complex ensembles in Rg-SASA space, together with the relationship between HADDOCK ranking and true structural quality. MD-induced relaxation and conformational exploration increased monomer diversity and broadened the accessible docking space. However, because of induced-fit rearrangements and limitations of the HADDOCK native score, a near-native conformation was not necessarily ranked highly even when it had been sampled (Supplementary Fig. S3F).

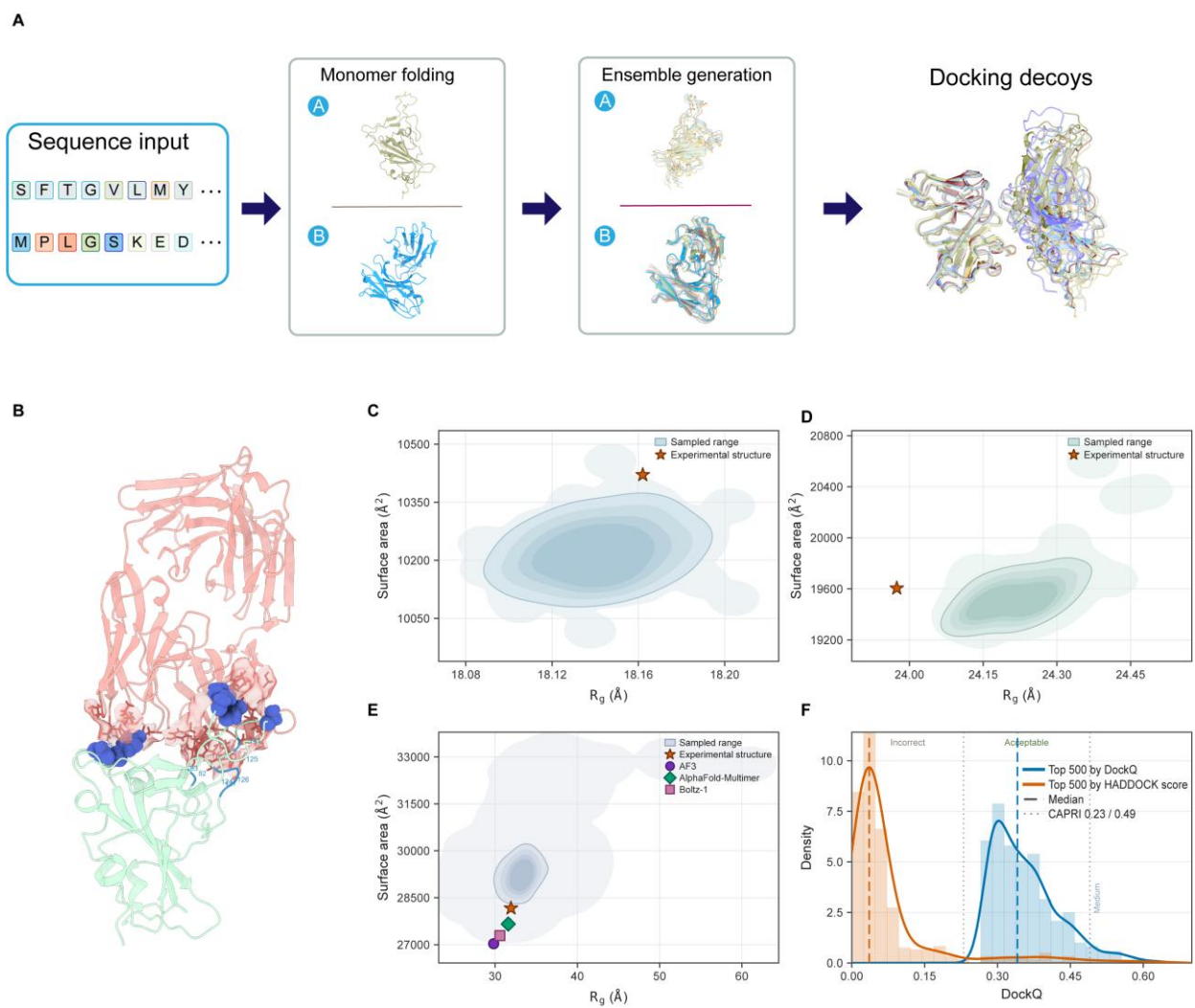

**Figure S3. MD-based expansion of monomer conformational ensembles and representative flexible-docking sampling.** (A) Workflow from sequence input and monomer folding to ensemble generation and docking-decoy construction. (B) The 7BEM antibody-antigen complex and interface anchor residues used to define ambiguous interaction restraints. (C, D) Rg-SASA distributions of the antigen and antibody monomer ensembles, respectively; stars mark the experimental structures. (E) Rg-SASA positions of complexes generated by AlphaFold3, AlphaFold-Multimer and Boltz-1; the background density shows the HADDOCK sampling region. (F) DockQ distributions for the 500 conformations with the highest DockQ values and the 500 conformations with the best HADDOCK scores. Dashed lines mark the medians, and dotted lines mark the DockQ thresholds of 0.23 and 0.49.

#### S3.4 Dataset composition and benchmark partitioning

The final training dataset contained 6,426 unique protein-complex targets listed in Supplementary Table 4. The external evaluation sets contained 201 AbSet blind-docking targets (Supplementary Table 5), 127 AbSet site-directed-docking targets (Supplementary Table 6) and

108 FoldBench targets (Supplementary Table 7). The internal antigen-antibody test set contained 102 targets (Supplementary Table 8).

#### S3.5 Target-level quality coverage of the conformation-augmented docking training data

For each of the 6,426 training targets, we selected the 500 highest finite DockQ values within [0, 1], ranked by DockQ rather than by the HADDOCK score. We then calculated the median, minimum and maximum DockQ values within this selected set. All training targets contributed 500 valid values. Figure S4 shows the corresponding CAPRI quality categories, and target-level values are available in Supplementary Table 9.

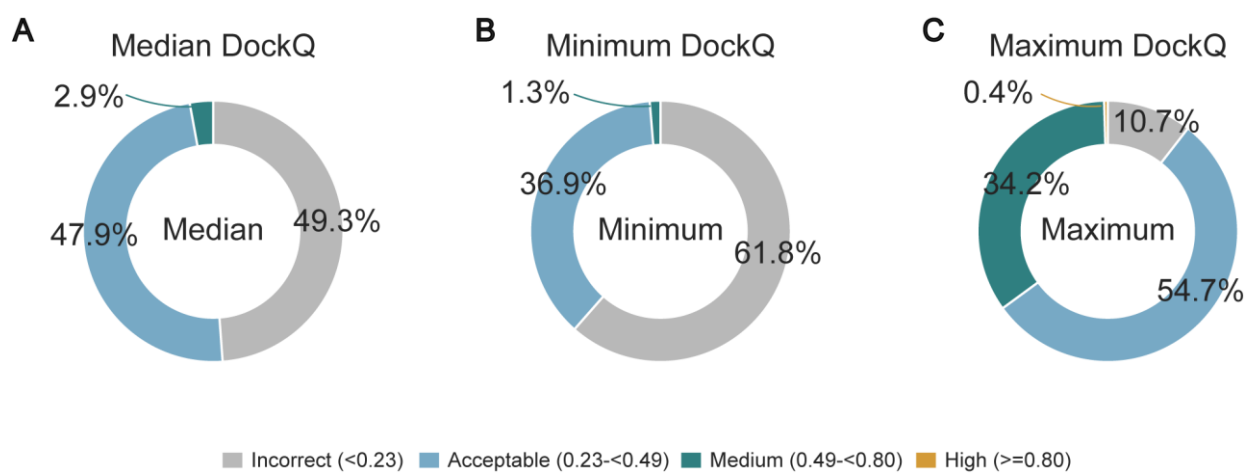

**Figure S4. Quality-category distributions of target-level top-500 DockQ summaries.** (A) Median DockQ among the 500 highest valid DockQ values per training target. (B) Minimum DockQ within the same selected set. (C) Maximum DockQ within the same selected set. Wedges show the percentage of the 6,426 training targets; every target contributed 500 finite DockQ values within [0, 1]. Categories are incorrect (DockQ < 0.23), acceptable (0.23 ≤ DockQ < 0.49), medium (0.49 ≤ DockQ < 0.80) and high (DockQ ≥ 0.80). The thresholds classify target-level summary statistics rather than all individual docking models. Source data are provided in Supplementary Table 9.

### S4. Quality-assessment model and training objective

#### S4.1 DockQ supervision target

DockQ was used as the primary supervision target. This metric integrates native-contact recovery, interface structural deviation and ligand displacement after receptor alignment:

$$S_i = \frac{1}{1 + \left(\frac{\text{IRMSD}}{1.5}\right)^2} \quad (\text{S4})$$

$$S_L = \frac{1}{1 + \left(\frac{\text{LRMSD}}{8.5}\right)^2} \quad (\text{S5})$$

$$\text{DockQ} = \frac{\text{Fnat} + S_i + S_L}{3} \quad (\text{S6})$$

Fnat denotes the fraction of native interface contacts recovered by a decoy. iRMSD is the root-mean-square deviation of the interface region, whereas LRMSD is the ligand deviation after receptor alignment. Si and SL map iRMSD and LRMSD to the interval 0-1, respectively. Higher DockQ indicates a higher-quality decoy. The model does not regress a single DockQ scalar; instead, it predicts Fnat, Si and SL separately and reconstructs DockQ as their mean. This design decomposes contact recovery, local interface geometry and global relative orientation into complementary supervision signals.

### S4.2 Input-structure standardization

Each decoy was first standardized as a two-protein complex, and the two binding partners to be evaluated were assigned as chains A and B. For multichain complexes, the interacting chain pair was selected before preprocessing. Only standard-protein ATOM records were used to build the graphs. Water, ligands, other HETATM records and hydrogen atoms were excluded, while carbon, nitrogen, oxygen, sulfur and other heavy atoms were retained. Residues lacking a Cα atom were marked invalid and excluded from graph construction. Unrecognized amino acids were mapped to an unknown residue class. Each valid residue retained its amino-acid type, chain identity, within-chain index, Cα coordinates, local coordinate frame, Cα-to-Cβ direction and interface label.

### S4.3 Cα residue representation

#### S4.3.1 Cα nodes

Each valid residue in the complete complex corresponds to one Cα node. The Cα graph represents global geometry, the relative positions of the chains and the global structural environment outside the interface. Each Cα node contains the following features:

**Table S1. Residue-node features in the global Cα graph.**

| Feature | Meaning |
| --- | --- |
| Amino-acid type | Distinguishes the 20 standard amino acids and an unknown class |
| Chain identity | Distinguishes chains A and B |
| Within-chain position | Sinusoidal positional encoding of the residue index |
| Local backbone conformation | Local dihedral-angle information derived from consecutive $C\alpha$ atoms |
| Local contact density | Number of $C\alpha$ neighbours within 10 Å |
| Terminal indicator | Marks the N- and C-terminal residues of each chain |
| Backbone direction | Local backbone-extension direction |
| Side-chain direction | Normalized $C\alpha$-to-$C\beta$ vector |

For glycine or residues lacking $C\beta$, an approximate side-chain direction derived from the local residue coordinate frame was used.

##### S4.3.2 Local $C\alpha$ coordinate frame

Each residue uses $C\alpha$ as the coordinate origin. The local x axis points from the current $C\alpha$ to the next $C\alpha$; together with the direction to the preceding $C\alpha$, this defines the local y and z axes through a cross product. The coordinate frame represents relative directions and rotations between residues.

##### S4.3.3 Edge construction for the $C\alpha$ graph

A spatial k-nearest-neighbour graph was constructed for each model conformation from three-dimensional $C\alpha$ distances. The 16 nearest $C\alpha$ neighbours were identified for each residue, and edges longer than 24 Å were removed. Directed edges allowed each residue to aggregate information from its spatial neighbours. Each edge encoded $C\alpha$-$C\alpha$ distance with radial basis functions, chain identity and within-chain sequence separation, as well as the position of residue j in the local coordinate frame of residue i, the position of residue i in the local coordinate frame of residue j and the relative rotation between the two frames. The global $C\alpha$ branch therefore captures both within-chain folding and the overall spatial relationship between chains.

##### S4.4 Identification of the protein interface and interface heavy atoms

$C\alpha$ distance alone can miss contacts formed by long side chains. Two residues may have distant $C\alpha$ atoms while their side chains form hydrogen bonds, salt bridges or hydrophobic contacts. We therefore identified the interface from the minimum distance between cross-chain non-hydrogen heavy atoms.

##### S4.4.1 Cross-chain residue-pair distance

For residue $r_i^A$ in chain A and residue $r_j^B$ in chain B, the residue-pair distance was defined as the minimum Euclidean distance between their non-hydrogen heavy atoms:

$$d_{ij}^{\text{res}} = \min_{a \in r_i^A, b \in r_j^B} \|x_a - x_b\|_2 \quad (\text{S7})$$

If the following condition was satisfied:

$$d_{ij}^{\text{res}} < 6.0 \text{ \AA} \quad (\text{S8})$$

both residues in the pair were labelled as interface residues. Specifically, a residue was labelled as an interface residue when any of its heavy atoms was within 6.0 Å of a heavy atom in the other chain.

##### S4.4.2 Handling sparse interfaces

In low-quality decoys, the two chains may be far apart, leaving few or no residue pairs within 6.0 Å. To retain the nearest contact region for model analysis, we used a fallback strategy. We first counted cross-chain residue pairs satisfying the 6.0 Å threshold. If fewer than five pairs were present, all cross-chain residue pairs were ranked by their minimum heavy-atom distance, the ten closest pairs were selected and the residues at both ends of each pair were labelled as candidate interface residues. This strategy prevents the interface branch from being empty for low-quality or clearly incorrect decoys.

##### S4.4.3 Interface-size limit

If the number of qualifying interface residues was too large, residues were ranked by their minimum distance to the other chain and the closest residues were retained, with a maximum of 100 interface residues per decoy. Not all atoms of an interface residue were required to lie within 6.0 Å of the other chain. Once any side-chain atom formed a close cross-chain contact, the residue was included in the interface heavy-atom graph together with its backbone and remaining side-chain atoms. The model therefore learned complete within-residue geometry rather than only direct contact pairs.

#### S4.5 Interface heavy-atom graph

##### S4.5.1 Atomic-node features

Each non-hydrogen heavy atom at the interface was represented as a node. Node features included element type (C, N, O, S or other), residue type, chain identity, backbone or side-chain identity and the encoded distance to the interface centre. The interface centre was defined as the mean coordinate of all interface heavy atoms in the current decoy. Distance to this centre distinguished core from peripheral interface atoms.

##### S4.5.2 Edge construction for the atomic graph

Edges were constructed between selected interface heavy atoms using a 4.5 Å distance cutoff:

$$(a, b) \in E_{\text{atom}}, a \neq b, \|x_a - x_b\|_2 < 4.5 \text{Å} \quad (\text{S9})$$

This rule permits edges within the same residue, between different residues in the same chain and between the two chains. Each edge encodes atomic distance, whether the atoms belong to the same residue and whether the edge crosses chains. The graph can therefore distinguish intramolecular geometry, local within-chain packing and genuine cross-chain interactions.

##### S4.5.3 Atom-to-residue aggregation

After atom-level geometric message passing, all heavy atoms within a residue were aggregated into a residue representation with attention weighting:

$$e_k = w^T \tanh(Wh_k) \quad (\text{S10})$$

$$\alpha_k = \frac{\exp(e_k)}{\sum_{l \in r} \exp(e_l)} \quad (\text{S11})$$

$$h_r^{\text{intf}} = \sum_{k \in r} \alpha_k h_k \quad (\text{S12})$$

The attention weights allow the model to identify the atoms that are most informative for interface quality in each decoy. Side-chain atoms forming cross-chain hydrogen bonds or salt bridges can receive higher weights than non-contact backbone atoms.

#### S4.6 Two-resolution structural encoding

The model contains two complementary structural branches:

**Table S2. Two branches of the two-resolution structural encoder.**

| Branch | Resolution | Main role |
| --- | --- | --- |
| Global Cα branch | Residue level | Captures global geometry, relative chain positions and long-range structural context |
| Interface heavy-atom branch | Atom level | Captures local interface contacts, side-chain packing and cross-chain geometry |

Both branches use residual geometric message passing. At each layer, messages are computed from neighbouring nodes and edge features, summed over neighbours and used to update the central node representation:

$$m_{ij} = \text{MLP}_{\text{edge}}[h_i, h_j, e_{ij}] \quad (\text{S13})$$

$$\bar{m}_i = \sum_j m_{ij} \quad (\text{S14})$$

$$h'_i = \text{LayerNorm}(h_i + \text{MLP}_{\text{node}}[h_i, \bar{m}_i]) \quad (\text{S15})$$

##### S4.7 Interface-dominant structural fusion

At interface residues, the interface heavy-atom branch is dominant and gated global Cα context is added:

$$g_{\text{ca}} = \sigma(W_{\text{ca}}[h_{\text{intf}}, h_{\text{ca}}, 1]) \quad (\text{S16})$$

$$h_{\text{core}} = h_{\text{intf}} + g_{\text{ca}} \odot h_{\text{ca}} \quad (\text{S17})$$

For non-interface residues, the interface heavy-atom representation is absent and only the attenuated gated Cα background is retained:

$$g_{\text{bg}} = \sigma(W_{\text{bg}}[h_{\text{ca}}, 0]) \quad (\text{S18})$$

$$h_{\text{core}} = g_{\text{bg}} \odot h_{\text{ca}} \quad (\text{S19})$$

##### S4.8 Late fusion of ESM sequence information

Each residue has a precomputed 1,280-dimensional ESM-2 representation. Sequence information does not enter the Cα or interface heavy-atom graphs; it is added only after the structural core representation has formed. ESM features are first projected to 128 dimensions. Interface residues then serve as queries for attention over the ESM key/value features of all valid residues in the complete complex. Interface residues can therefore access both local sequence features and long-range sequence context. ESM updates in each cross-attention module are controlled by a channel-wise gate:

$$g_{\text{esm}} = \sigma(W_g[q, q_{\text{esm}}, w_{\text{intf}}]) \quad (\text{S20})$$

$$q' = \text{LayerNorm}(q + g_{\text{esm}} \odot s \odot \text{MHA}(q, \text{kv}, \text{kv})) \quad (\text{S21})$$

where $s$ is a learnable LayerScale parameter.

##### S4.9 Sample-level dynamic readout

The model aggregates three representations over interface residues: the interface heavy-atom branch, the global Cα features at interface positions and the ESM-enhanced interface features. Attention pooling produces a 128-dimensional decoy representation from each branch, after which dynamic branch weights are calculated from the three vectors for the current decoy:

$$[\alpha_{\text{intf}}, \alpha_{\text{ca}}, \alpha_{\text{esm}}] = \text{softmax}(\text{MLP}[z_{\text{intf}}, z_{\text{ca}}, z_{\text{esm}}]) \quad (\text{S22})$$

$$z = \alpha_{\text{intf}} z_{\text{intf}} + \alpha_{\text{ca}} z_{\text{ca}} + \alpha_{\text{esm}} z_{\text{esm}} \quad (\text{S23})$$

##### S4.10 DockQ component prediction

The dynamic fused representation is projected to 512 dimensions and then fed into three independent prediction heads. All three heads share an identical architecture: the 512-dimensional input is first transformed by a hidden layer of 64 units and then mapped to a single scalar, followed by a sigmoid activation. The three heads respectively output the predicted Fnat, Si, and SL scores.

All three components are constrained to 0-1, and the predicted DockQ is reconstructed as:

$$\text{DockQ} = \frac{\hat{\text{Fnat}} + \hat{S}_i + \hat{S}_L}{3} \quad (\text{S24})$$

##### S4.11 Sample selection at each training step

###### S4.11.1 Grouping decoys by target

All decoys were first grouped by protein-complex target. Ranking was learned only within the same target, without comparing DockQ values across proteins. Using DockQ = 0.23 as the boundary, decoys were divided into a better group (DockQ ≥ 0.23) and a poorer group (DockQ < 0.23). DockQ = 0.23 is the lower boundary of acceptable quality.

###### S4.11.2 Composition of training samples

Each training item first samples one target and then 16 decoy conformations from that target.

A target was selected at random. To account for quality stratification, two decoys were sampled from its better group and 14 from its poorer group. The 16 decoys were pooled and shuffled. Regression loss was calculated for every decoy, and ranking pairs were constructed within the set. If one group contained too few decoys, all available samples from that group were used before sampling with replacement. If a target contained only one group, all 16 samples were drawn from the available group.

##### S4.12 Piecewise DockQ sample weights

The training data were typically dominated by low-quality decoys. Equal weighting would allow the numerous incorrect conformations to dominate the gradient and bias fitting toward low DockQ values, reducing attention to the medium- and high-quality conformations most important for Top-k ranking. Each sample therefore received a quality weight based on its true DockQ:

**Table S3. Sample weights defined by DockQ interval.**

| DockQ interval | Quality class | Sample weight |
| --- | --- | --- |
| < 0.23 | Incorrect | 1.0 |
| 0.23-0.49 | Acceptable | 2.0 |
| 0.49-0.80 | Medium | 3.0 |
| $\geq 0.80$ | High | 4.0 |

DockQ values exactly equal to 0.23, 0.49 or 0.80 were assigned to the higher-weight interval. The weights did not change the labels but adjusted the relative contribution of samples to the gradient. The weighted loss was normalized by the sum of weights:

$$L = \frac{\sum_i w_i \ell_i}{\sum_i w_i} \quad (\text{S25})$$

#### S4.13 Multi-component regression loss

##### S4.13.1 Smooth L1 loss

Fnat, Si, SL and reconstructed DockQ were all optimized with Smooth L1 loss. For prediction error $e = \hat{y} - y$, $\beta$ was set to 0.1:

$$\ell_{\text{smoothL1}}(e) = \begin{cases} \frac{e^2}{2\beta} & |e| < \beta \\ |e| - \frac{\beta}{2} & |e| \geq \beta \end{cases} \quad (\text{S26})$$

The quadratic penalty for small errors promotes accurate fitting, whereas the approximately linear penalty for larger errors reduces the influence of extreme outliers on training stability.

##### S4.13.2 Four regression terms

The regression loss was defined as:

$$L_{\text{reg}} = 1.0L_{\text{Fnat}} + 1.0L_{\text{Si}} + 1.0L_{\text{SL}} + 0.3L_{\text{DockQ}} \quad (\text{S27})$$

Each loss term $L$ used the piecewise DockQ sample weights above. Fnat measures contact recovery and helps distinguish acceptable from incorrect conformations. Si and SL constrain local interface geometry and the global ligand position, respectively. Although DockQ is reconstructed from these components, the separate DockQ term directly constrains the final ranking score toward the true DockQ.

#### S4.14 Target-wise pairwise ranking loss

Numerical regression alone does not guarantee that high-quality decoys from the same target will rank above low-quality decoys. An explicit ranking loss was therefore added to the regression objective.

##### S4.14.1 Positive and negative samples with a buffer interval

Ranking pairs used the following thresholds: positive samples had DockQ > 0.23 and clear negative samples had DockQ < 0.21. Samples with DockQ between 0.21 and 0.23 formed a boundary buffer and were excluded from positive-negative pairing. This buffer reduces unstable supervision near the 0.23 boundary caused by label noise or small structural differences.

##### S4.14.2 Positive-negative pairs

For any positive sample $i$ and clear negative sample $j$ within the same target, the following condition was imposed:

$$\hat{D}_i > \hat{D}_j \quad (\text{S28})$$

Here, $\hat{D}$ denotes the predicted DockQ.

##### S4.14.3 Ranking within the positive set

For two positive samples whose true DockQ values differed by at least 0.03, the higher-quality sample was also required to receive the higher predicted score:

$$D_i - D_j \geq 0.03 \Rightarrow \hat{D}_i > \hat{D}_j \quad (\text{S29})$$

The model therefore learns both to separate acceptable from incorrect conformations and to order acceptable, medium and high-quality conformations more finely.

##### S4.14.4 Pairwise logistic loss

For each ordered pair $(i, j)$, when $i$ should rank ahead of $j$, the following loss was used:

$$\ell_{\text{rank}}(i, j) = \text{softplus} \left( -\frac{\hat{D}_i - \hat{D}_j}{\tau} \right) \quad (\text{S30})$$

The temperature parameter $\tau$ was set to 0.1, without an additional explicit reward term. When the ordering is correct, the loss approaches zero as the score difference increases; when the ordering is reversed, the loss increases rapidly.

##### S4.14.5 Pair weights

Each pair was weighted by the quality weight of the higher-DockQ sample that should rank first. Thus, incorrect ranking of high-quality conformations received the strongest penalty, followed by incorrect ranking of medium-quality conformations.

##### S4.15 Total loss

The final training objective was:

$$L_{\text{total}} = L_{\text{reg}} + 0.2L_{\text{rank}} \quad (\text{S31})$$

Expanding this objective gives:

$$L_{\text{total}} = 1.0L_{\text{Fnat}} + 1.0L_{\text{Si}} + 1.0L_{\text{SL}} + 0.3L_{\text{DockQ}} + 0.2L_{\text{rank}} \quad (\text{S32})$$

The regression loss estimates absolute conformation quality, whereas the ranking loss constrains the relative order of decoys within the same target. Together, they produce physically interpretable DockQ components while optimizing the Top-k reranking required in practical structure prediction.

### S5. FACET ablation experiments

To quantify the contribution of each FACET module, we evaluated the full model and four ablation variants on the same protein-level five-fold cross-validation split. Each variant removed or simplified one component: the global Cα equivariant graph neural network (EGNN), the interface heavy-atom EGNN, the structural fusion gate or the late ESM fusion. All models used the same data partitions and validation sets to limit differences unrelated to architecture.

Within each protein complex, and reflecting the intended ranking use case, we calculated Spearman and Kendall rank correlations across all candidate conformations and then summarized the results over protein complexes in five mutually exclusive test folds. Each protein was weighted equally, regardless of the number of candidate conformations. The full model and each ablation were compared on paired protein sets with two-sided Wilcoxon signed-rank tests; within each prediction-target/metric combination, P values for the four ablation comparisons were corrected with the Holm method.

Across 32 prediction combinations, the full model had a higher mean in 30 cases, and 28 remained significant after Holm correction. Removing the global Cα EGNN or late ESM fusion significantly reduced all eight metrics. Removing the interface heavy-atom EGNN reduced six of eight metrics significantly, whereas the two DockQ differences were not significant. Removing the structural fusion gate significantly reduced the six comparisons for DockQ, Si and SL, but the ablation was slightly higher for Fnat Spearman and Kendall correlations. Late ESM fusion made the largest and most consistent contribution: removing it reduced Spearman correlations by 0.0189-0.0690 and Kendall correlations by 0.0160-0.0505, with all eight comparisons remaining significant after Holm correction. Overall, the protein-level paired evaluation supports a broad advantage for the full architecture, with the strongest evidence for late ESM fusion. Supplementary Figs. S5 and S6 show the ablation results and statistical validation.

These results indicate that the global and local structural encoders generally provide complementary information. The exception observed after removing the structural fusion gate for

Fnat suggests that contact recovery and global conformational quality may respond differently to global $C\alpha$ and interface-atom features, but this does not offset the consistent gains for DockQ, Si and SL.

**A**

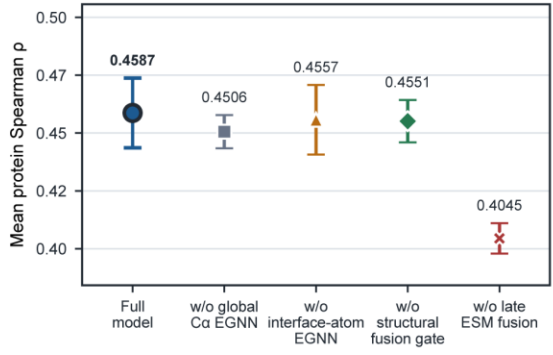

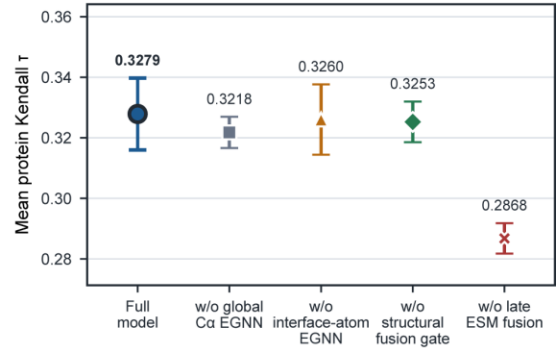

**B**

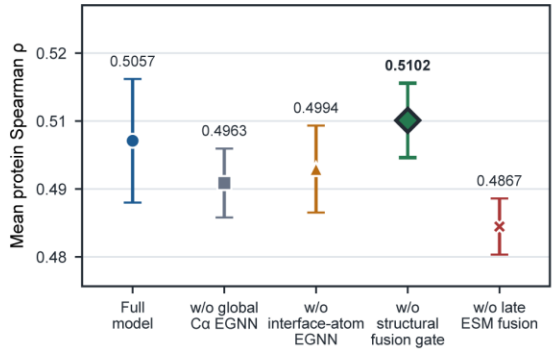

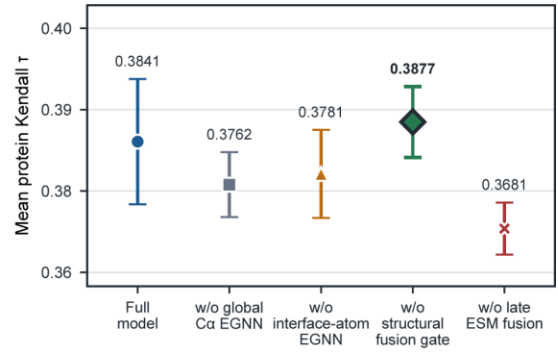

**C**

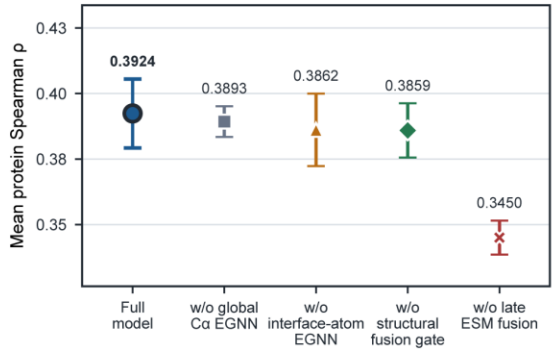

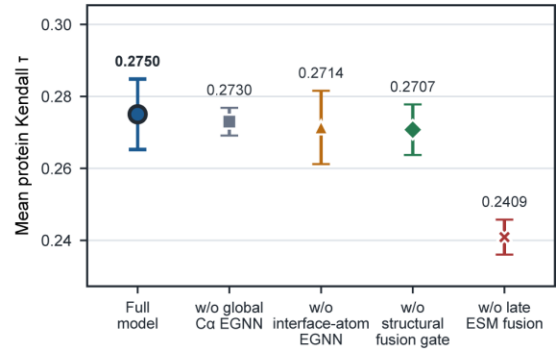

**D**

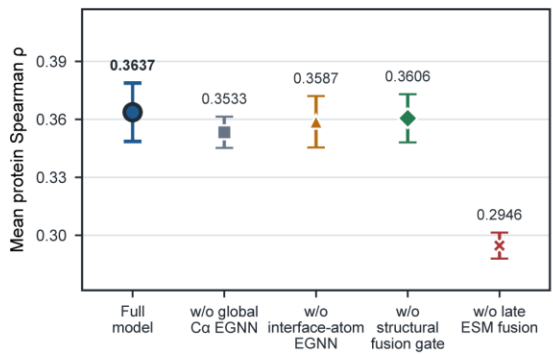

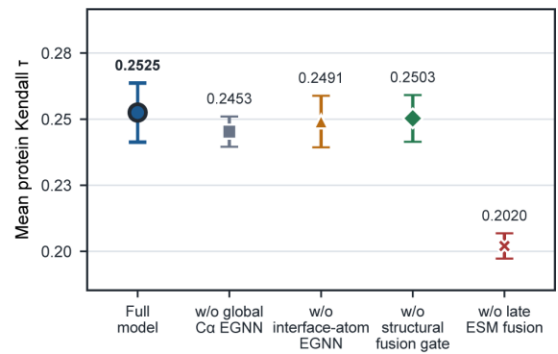

**Figure S5. Protein-level rank correlations for the full FACET model and four ablation variants. (A-D)** Results for DockQ, Fnatt, Si and SL, respectively. Within each panel, the left plot shows mean within-protein Spearman $\rho$ and the right plot shows mean within-protein Kendall $\tau$. Points and error bars show the mean ± standard error across five target-level cross-validation folds.

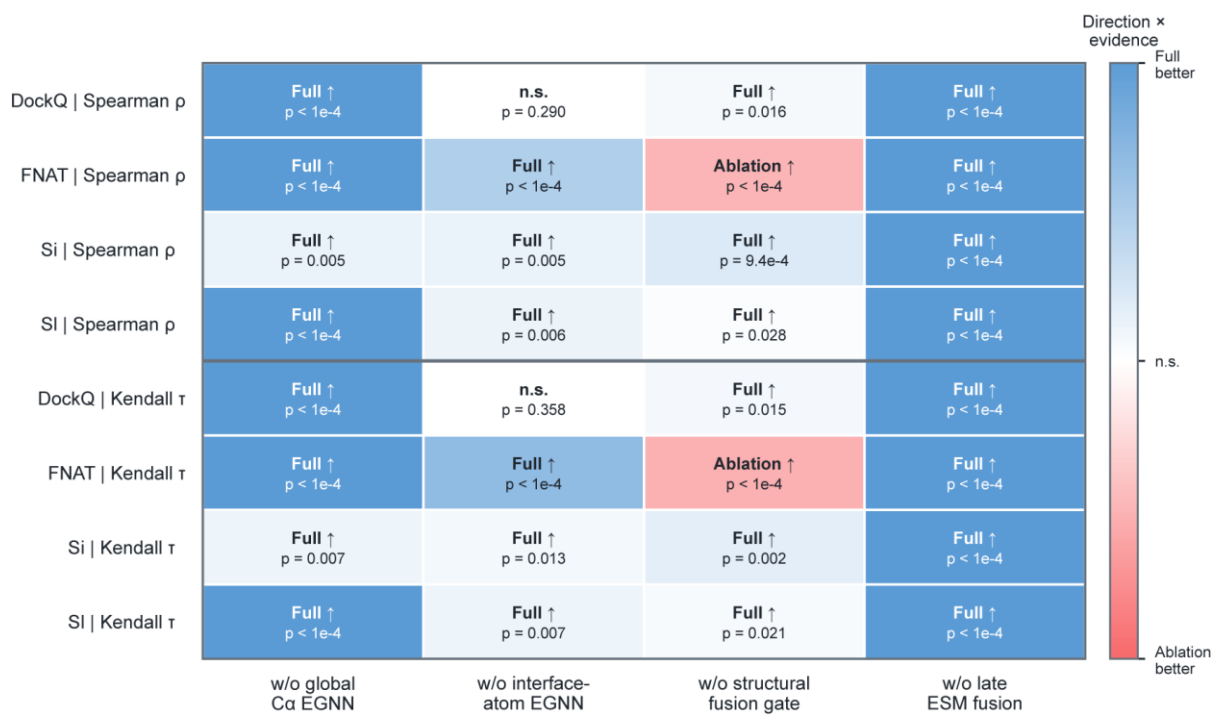

**Figure S6. Direction and significance of paired protein-level comparisons between the full FACET model and four ablation variants.** Each cell reports the direction of the mean paired difference and the Holm-adjusted p value from a two-sided Wilcoxon signed-rank test. Adjustment was performed across the four ablations for each prediction-target and correlation-metric combination. Blue indicates a significantly higher correlation for the full model, red indicates a significantly higher correlation for the ablation, and white indicates no significant difference.

### S6. FACET compared with graph-neural-network baselines

As a complementary evaluation of the FACET implementation, we compared it with four graph-neural-network architectures: HGT, GAT, GCN and DGN. Data were split by protein target into five-fold cross-validation partitions. Global Spearman and Kendall rank correlations, mean absolute error (MAE) and root-mean-square error (RMSE) were used to evaluate candidate ranking and DockQ regression performance (Supplementary Fig. S7).

FACET achieved global Spearman and Kendall correlations of 0.6700 and 0.4862, respectively, compared with 0.5885 and 0.4186 for HGT. FACET also achieved the lowest MAE

and RMSE, 0.0849 and 0.1177, respectively, compared with 0.0982 and 0.1301 for HGT and 0.1110 and 0.1408 for DGN. Among the evaluated architectures, FACET therefore provided the best overall balance between candidate-ranking correlation and DockQ regression accuracy.

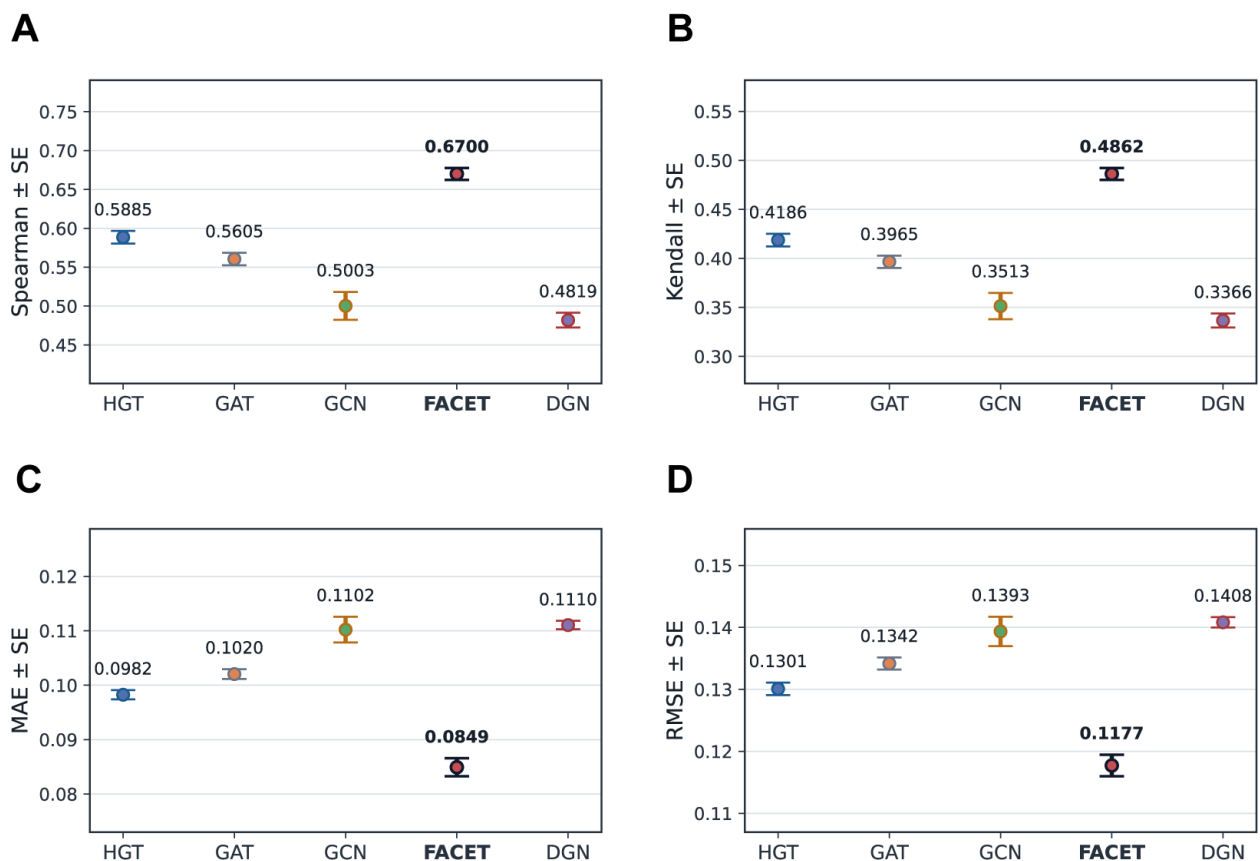

**Figure S7. DockQ prediction performance of FACET and the graph-neural-network baselines HGT, GAT, GCN and DGN.** (A) Global Spearman correlation. (B) Global Kendall correlation. (C) Mean absolute error (MAE). (D) Root-mean-square error (RMSE). Points and error bars show the mean ± standard error across five target-level cross-validation folds.

### S7. Supplementary reranking cases

We further examined representative targets in which FACET reranked candidates generated by Boltz-1 or AlphaFold-Multimer. For the targets shown, FACET recovered at least one acceptable model (DockQ $\geq 0.23$) within the top 10 candidates, whereas the corresponding native confidence score did not. Supplementary Figs. S8-S13 show the Top-1 structures selected from the same candidate sets.

### 7C01 | Rank 1 structural comparison

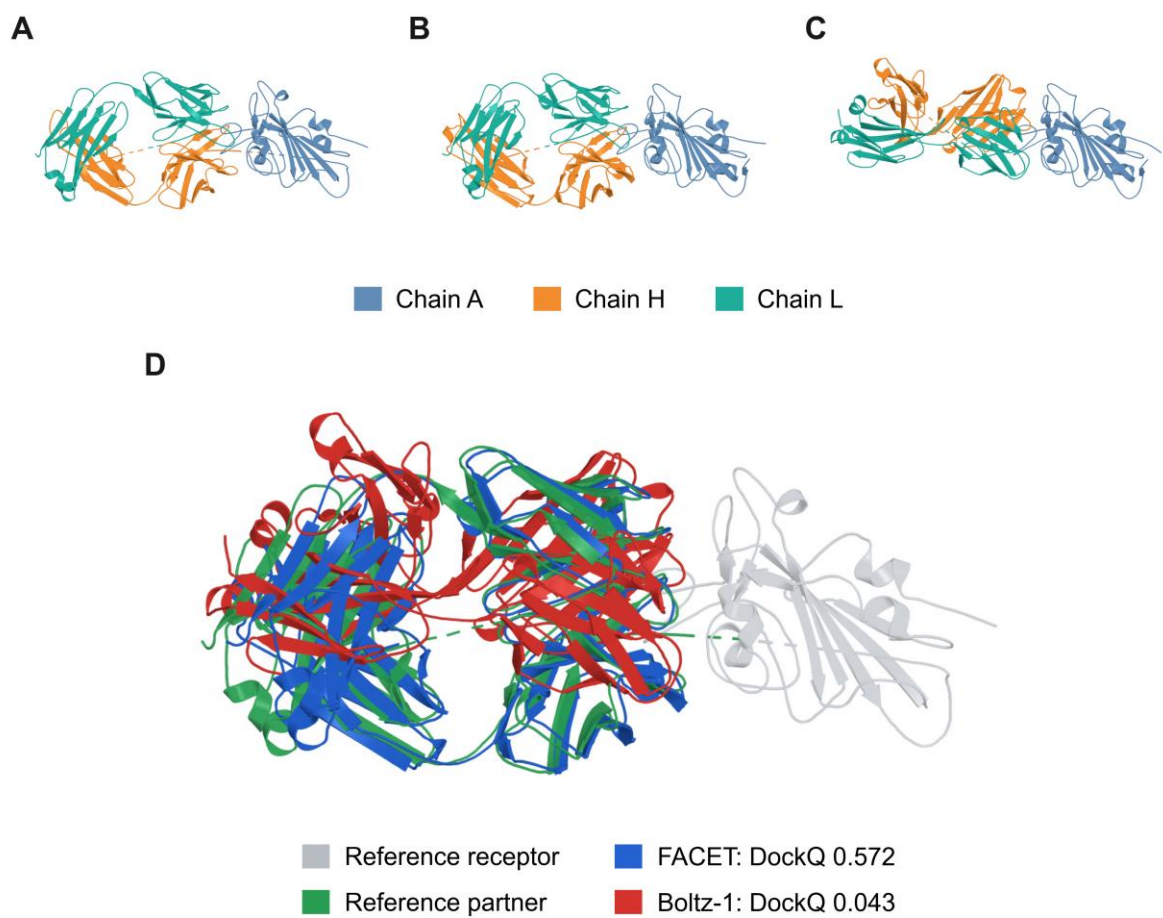

**Figure S8. Rank-1 structural comparison for PDB 7C01.** (A) Experimental reference complex, with chains A, H, and L shown in blue, orange, and teal. (B,C) Rank-1 predictions selected by FACET (DockQ = 0.572) and the Boltz-1 confidence score (DockQ = 0.043), respectively. (D) Receptor-aligned overlay; the reference receptor and partner are gray and green, and the FACET- and Boltz-1-selected models are blue and red.

### 7NEG | Rank 1 structural comparison

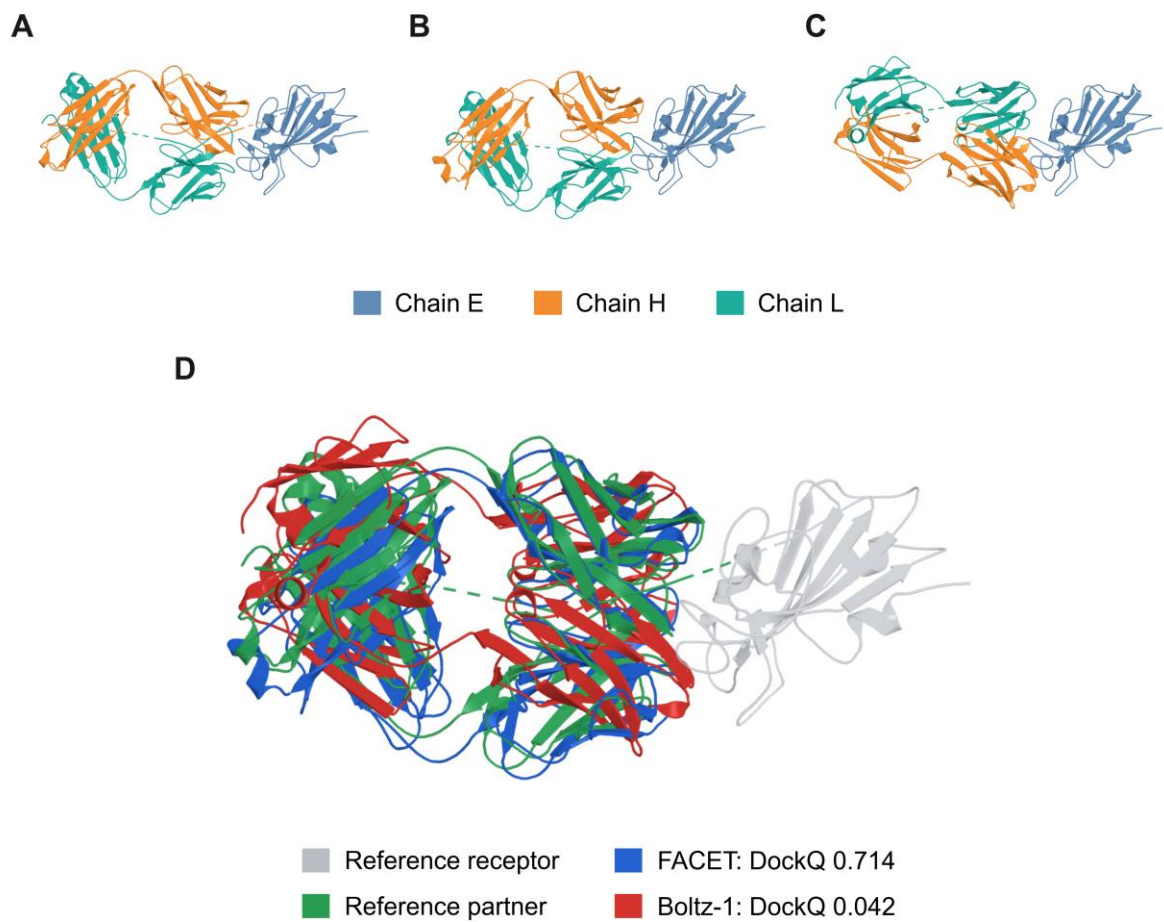

**Figure S9. Rank-1 structural comparison for PDB 7NEG.** (A) Experimental reference complex, with chains E, H, and L shown in blue, orange, and teal. (B,C) Rank-1 predictions selected by FACET (DockQ = 0.714) and the Boltz-1 confidence score (DockQ = 0.042), respectively. (D) Receptor-aligned overlay; the reference receptor and partner are gray and green, and the FACET- and Boltz-1-selected models are blue and red.

### 8BH5 | Rank 1 structural comparison

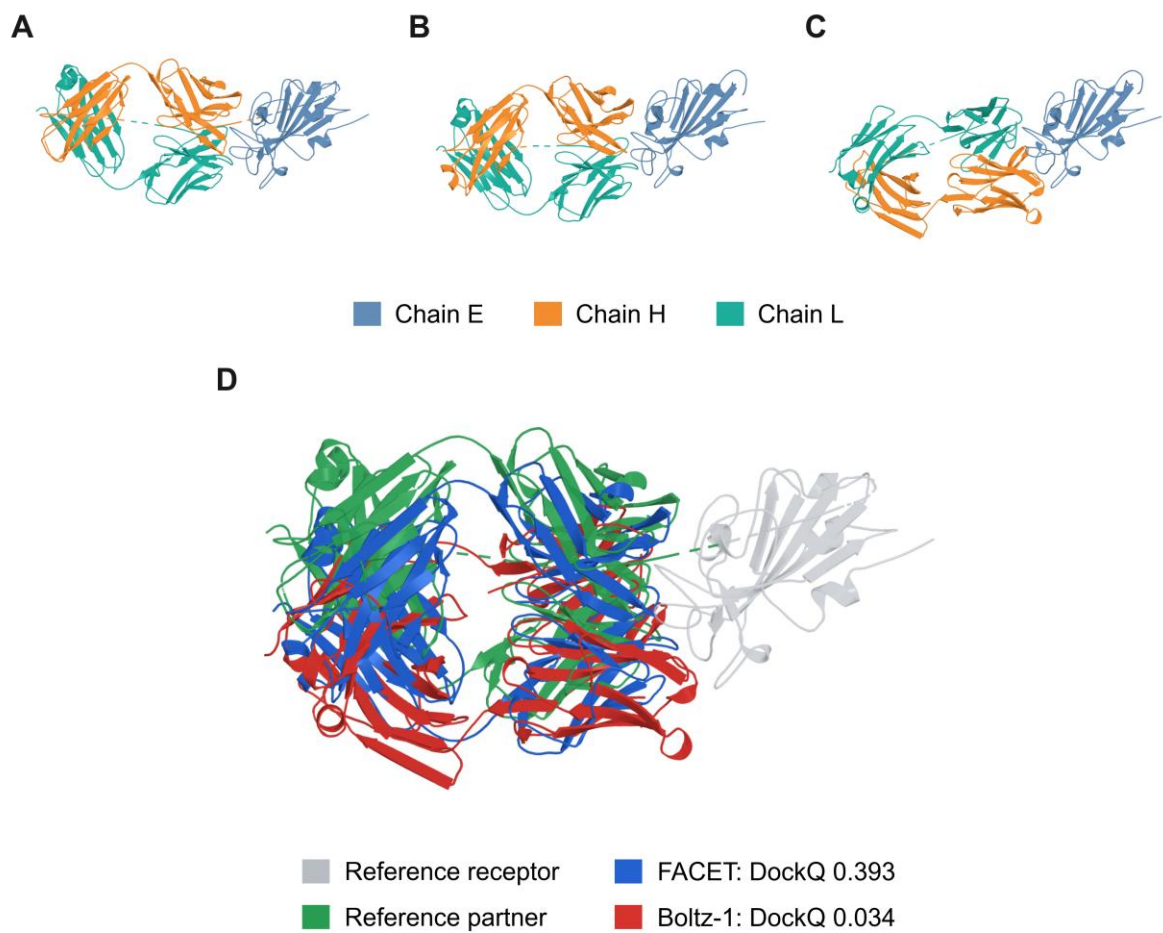

**Figure S10. Rank-1 structural comparison for PDB 8BH5.** (A) Experimental reference complex, with chains E, H, and L shown in blue, orange, and teal. (B,C) Rank-1 predictions selected by FACET (DockQ = 0.393) and the Boltz-1 confidence score (DockQ = 0.034), respectively. (D) Receptor-aligned overlay; the reference receptor and partner are gray and green, and the FACET- and Boltz-1-selected models are blue and red.

#### 5GJS | Rank 1 structural comparison

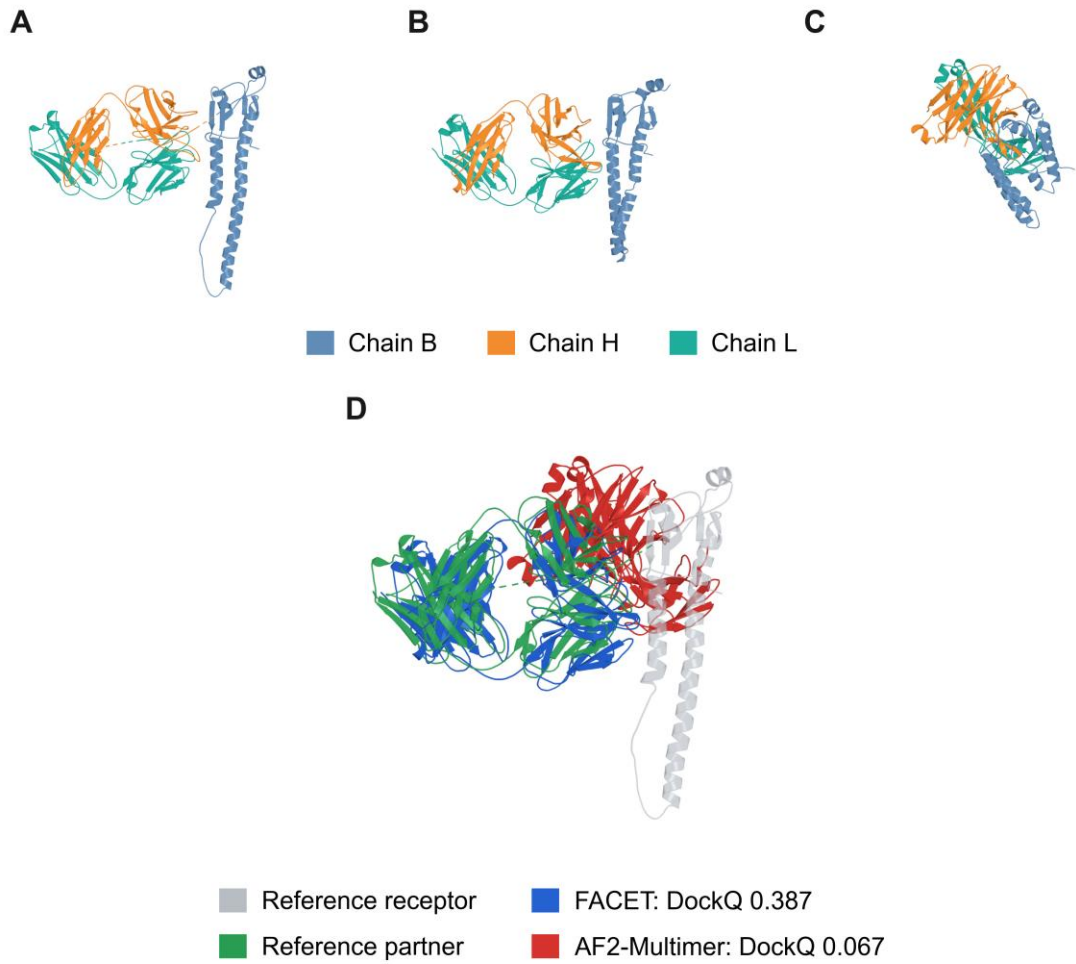

**Figure S11. Rank-1 structural comparison for PDB 5GJS.** (A) Experimental reference complex, with chains B, H and L shown in blue, orange and teal, respectively. (B,C) Rank-1 predictions selected by FACET (DockQ = 0.387) and AF2-Multimer (DockQ = 0.067), respectively. (D) Receptor-aligned overlay; the reference receptor and partner are gray and green, and the FACET- and AF2-Multimer-selected models are blue and red.

### 7EK0 | Rank 1 structural comparison

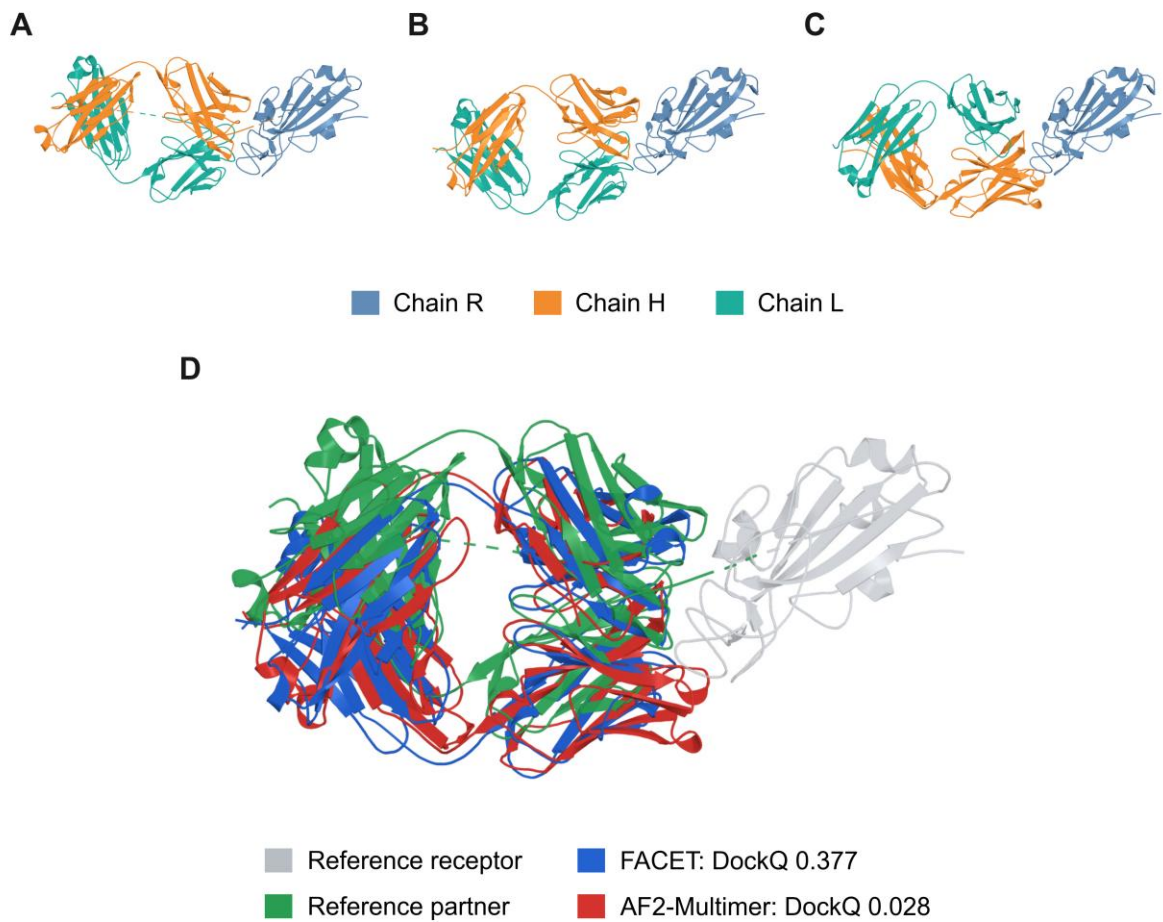

**Figure S12. Rank-1 structural comparison for PDB 7EK0.** (A) Experimental reference complex, with chains R, H and L shown in blue, orange and teal, respectively. (B,C) Rank-1 predictions selected by FACET (DockQ = 0.377) and AF2-Multimer (DockQ = 0.028), respectively. (D) Receptor-aligned overlay; the reference receptor and partner are gray and green, and the FACET- and AF2-Multimer-selected models are blue and red.

### 7KMI | Rank 1 structural comparison

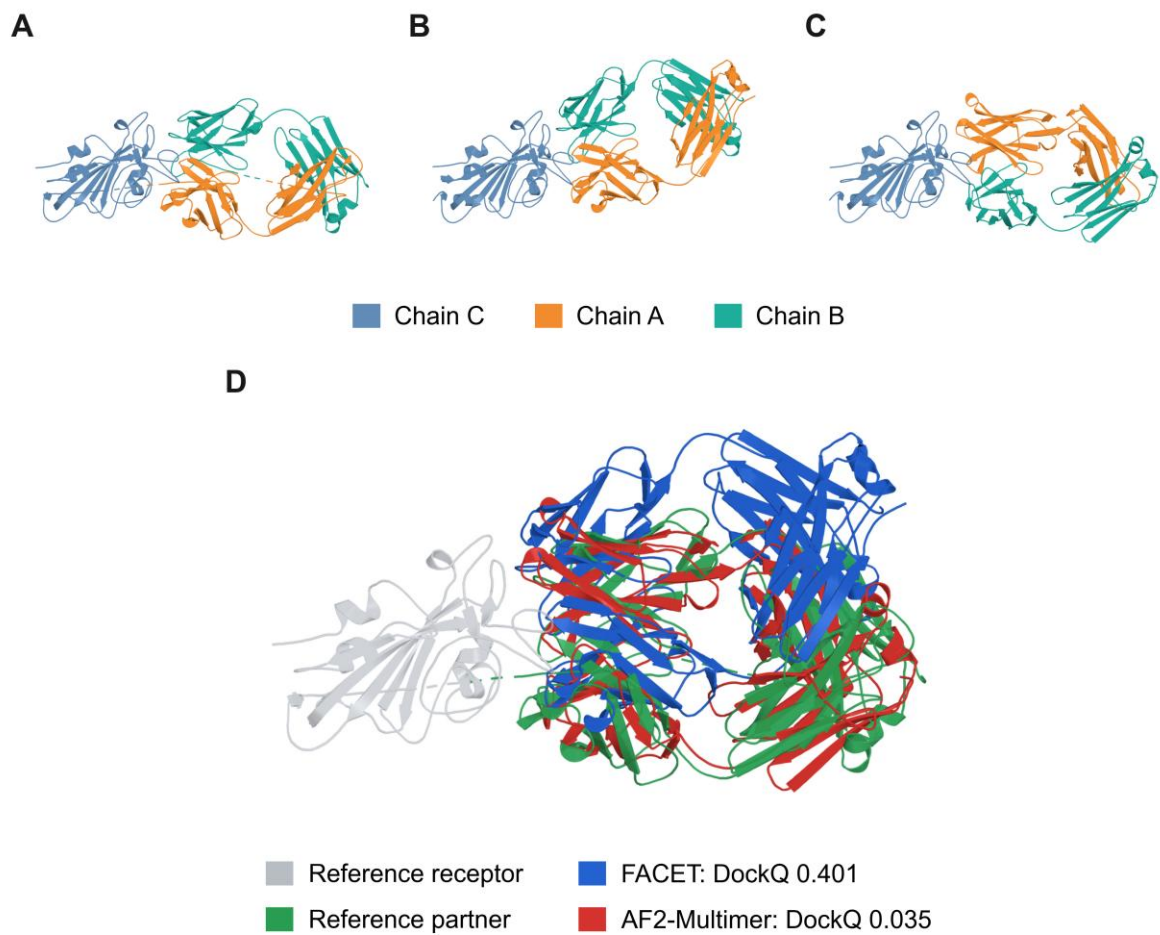

**Figure S13. Rank-1 structural comparison for PDB 7KMI.** (A) Experimental reference complex, with chains C, A and B shown in blue, orange and teal, respectively. (B,C) Rank-1 predictions selected by FACET (DockQ = 0.401) and AF2-Multimer (DockQ = 0.035), respectively. (D) Receptor-aligned overlay; the reference receptor and partner are gray and green, and the FACET- and AF2-Multimer-selected models are blue and red.
